# Midkine expression by stem-like tumor cells drives persistence to mTOR inhibition and rewires the microenvironment toward an immune-suppressive state

**DOI:** 10.1101/2022.06.01.494055

**Authors:** Yan Tang, David J. Kwiatkowski, Elizabeth P. Henske

**Author notes:** Corresponding Authors: David J. Kwiatkowski, M.D, Ph.D. Elizabeth P. Henske, M.D 20 Shattuck Street, Division of Pulmonary Medicine and Critical Care Medicine, Brigham and Women’s Hospital, Harvard Medical School, Boston, MA 02115, USA.

## Abstract

mTORC1 regulates cell metabolism to enable cell proliferation, and is hyperactive in multiple cancer types^1, 2^. Here, we performed integrative analysis of single cell transcriptomic profiling, paired T cell receptor (TCR) sequencing, and spatial transcriptomic profiling on Tuberous Sclerosis Complex (TSC) associated tumors with mTORC1 hyperactivity, and identified a stem-like tumor cell state (SLS) linked to T cell dysfunction via tumor-modulated immunosuppressive macrophages. Rapamycin and its derivatives (rapalogs) are the primary treatments for TSC tumors, and the stem-like tumor cells showed rapamycin resistance *in vitro*, reminiscent of the cytostatic effects of these drugs in patients. The pro-angiogenic factor midkine (MDK) was highly expressed by the SLS population, and associated with enrichment of endothelial cells in SLS-dominant samples. Inhibition of MDK showed synergistic benefit with rapamycin in reducing the growth of TSC cell lines *in vitro* and *in vivo*. In aggregate, this study suggests an autocrine rapamycin resistance mechanism and a paracrine tumor survival mechanism via immune suppression adopted by the stem-like state tumor cells with mTORC1 hyperactivity. We also provide a comprehensive resource to advance the understanding of TSC and potentially other mTORC1-hyperactive tumors.

## Introduction

Tuberous Sclerosis Complex (TSC) is an autosomal dominant disease with an incidence of 1:6000 births. TSC is caused by loss-of-function mutations in the tumor suppressor genes *TSC1* and *TSC2*^3^. Second hit loss of the remaining wild-type copy of *TSC1* or *TSC2* leads to hyperactive mTORC1, and drives tumor growth in multiple organs. ^3^. Angiomyolipoma (AML) and lymphangioleiomyomatosis (LAM) are common and related manifestations of TSC that can lead to renal and pulmonary insufficiency, respectively^4, 5^. AML and LAM also occur sporadically in patients without TSC^3–5^. The mTORC1 inhibitors sirolimus (rapamycin) and everolimus (Afinitor) are closely related compounds termed rapalogs, and are FDA-approved for the therapy of LAM and AML, respectively. Rapalogs induce a modest response in most patients with a median 50% volume reduction of AML^6^ and stabilization of lung function in LAM for at least 12 months^7^, with recurrent tumor growth and lung function decline after treatment cessation. Therapeutic strategies that eliminate, rather than suppress, tumor cells in TSC, are urgently needed.

Prior efforts to characterize TSC tumors using bulk RNA-Sequencing (RNA-Seq) has advanced our understanding of the unique transcriptional programs of TSC tumors^8^, including the important role of Melanocyte Inducing Transcription Factor (MITF)^9^, but were limited in the ability to reveal tumor cell heterogeneity, or interaction between tumor and microenvironment^8^. In contrast, single cell RNA-Sequencing (scRNA-Seq) enables comprehensive investigation of heterogeneity of tumor and microenvironment cells and global mapping of molecular interactions among cell types. Two recent single cell studies on LAM lungs have yielded important insight into the cellular origin of LAM cells and revealed alveolar epithelial remodeling by LAM cells^10, 11^. However, these studies were limited by the small number of LAM cells identified (< 200 LAM cells).

Tumor cell heterogeneity and plasticity is increasingly recognized as an important and common aspect of tumor biology. The occurrence of multiple cell states in tumors and plasticity of inter-conversion of cell states likely contributes to therapeutic resistance^12^. In AML, three different cell types represent the neoplastic process (fat, muscle, and vessels)^13^. Cellular heterogeneity is evident in both AML and LAM, but the precise components of this heterogeneity, how the different cellular elements inter-relate, and how each element responds to therapy are unexplored. In addition, aberrant vascular hypertrophy is also typical of AML^13^, and may contribute to an hypoxic tumor microenvironment. Tumor cells can acquire stemness and dormancy due to hypoxic conditions, and become stress and therapy resistant^14^.

Emerging data suggest that the immune system plays a key role in the pathogenesis and potentially the therapy of LAM and AML. Natural killer cells are enriched and activated in LAM^15, 16^. Evidence of T cell infiltration and exhaustion have been observed in human AML and LAM and in mouse models, and there is clear benefit of immunotherapy in mouse models of TSC and LAM^17, 18^. This T cell infiltration and dysfunction are unexpected since AML have a very low neoantigen burden^19^. Macrophage infiltration was also observed in renal AML^20^, hepatic AML^21^ and TSC skin tumors^22^. Despite these advances in understanding the immune microenvironment of LAM and AML, a comprehensive analysis has not been possible. In addition, the identification of molecular interactions between AML/LAM tumor cells and other cell types in the microenvironment has not previously been possible.

To address these points, we interrogated the tumor microenvironment of AML and LAM. Single cell profiling of 5 LAM specimens, 6 AML and 4 matched normal kidneys revealed two distinct cell states in AML/LAM cells: a stem-like state (SLS) and an inflammatory state (IS). SLS tumor cells exhibited high stemness and dormancy marker expression, and showed rapamycin resistance in primary angiomyolipoma-derived cultures. *MDK* was highly expressed specifically in SLS cells, and MDK inhibitor treatment enhanced the therapeutic effect of rapamycin in patient-derived TSC2-deficient AML cells *in vitro* and *in vivo*. Integrative analysis of single cell data and spatial transcriptomic profiling of these tumors further revealed a modulatory axis from SLS tumor cells to suppressive *TREM2*+/*TYROBP*+ macrophages, leading to T cell dysfunction. Concurrent single cell T cell receptor sequencing (scTCR-Seq) analysis demonstrated a substantial suppression of clonal expansion and T cell RNA velocity in SLS-dominant tumors compared to IS-dominant tumors. In contrast, inflammatory state (IS) tumor cells with low *MDK* expression showed high expression of cytokines and were enriched with immune regulatory pathways. Substantial T cell clonal expansion with elevated cytotoxic programs was observed in IS-dominant tumors compared with SLS-dominant tumors. Taken together, these data reveal differential immune remodeling by previously unrecognized distinct cells states in mTORC1-hyperactive tumors, and provide a rationale for precision immunotherapy in TSC.

## Results

### Single cell analysis of angiomyolipomas (AML) and lymphangioleiomyomatosis (LAM)

AML and LAM are hallmark manifestations of TSC^3^, and are also seen sporadically in patients without TSC. Six renal AML tumors and four matched normal tissues (Supplementary data 1) obtained at the time of tumor resection were assessed with scRNA-Seq and paired scTCR-Seq using the 10x Chromium single cell 5’ chemistry (Fig. 1a). Five LAM lungs (Supplementary data 1) obtained at lung transplantation were also analyzed with scRNA-Seq. After filtering out low-quality cells, a total of 108,071 cells from the AML and 33,136 cells from the matched normal kidneys were analyzed; 42,202 cells were analyzed from the LAM lung samples. Pathological images for the AML/LAM samples are provided in Supplementary Fig. 1.

**Fig. 1.**
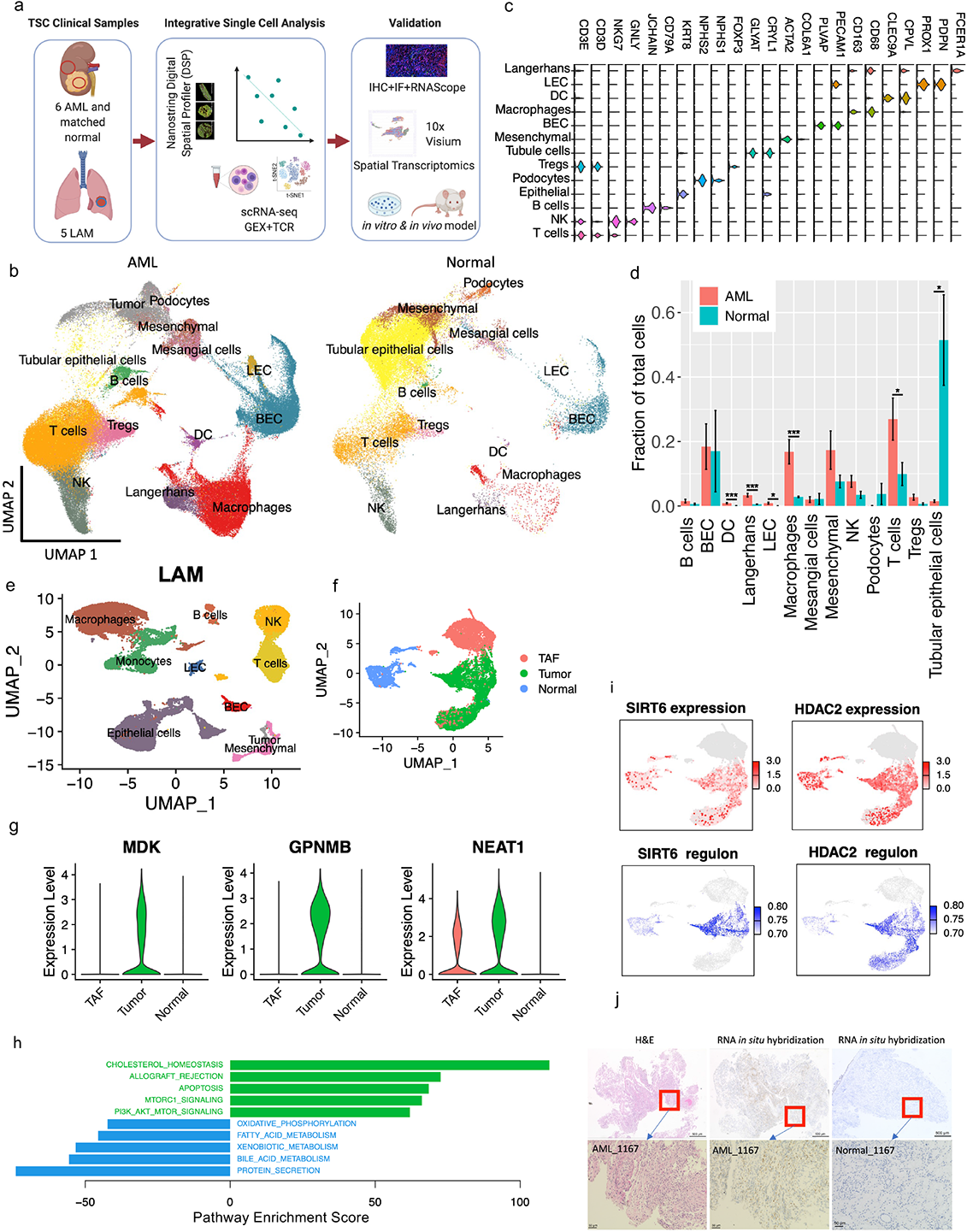
Single-cell atlas of angiomyolipoma (AML) and lymphangioleiomyomatosis (LAM). a. Workflow showing samples collected and integrative analysis of scRNA-Seq, paired scTCR-Seq and spatial transcriptomics, followed by *in vitro* and *in vivo* mechanistic studies. b. Uniform Manifold Approximation and Projection (UMAP) plots of major cell types identified in six AML tumors (left) and four matched normal kidneys (right). LEC: lymphatic endothelial cells; BEC: blood endothelial cells; Tregs: regulatory T cells; NK: natural killer cells; DC: dendritic cells. c. Violin plots of marker genes of each cell type. The y axis represents the normalized gene expression value. d. Quantification of fractional representation of cell types in tumors and matched normal tissues. Standard errors are shown for each group. *p < 0.05, **p<0.01, ***p<0.001, t-test. e. UMAP plot showing major cell types identified in five LAM lungs. LEC: lymphatic endothelial cells; BEC: blood endothelial cells; NK: natural killer cells. f. Re-clustering of mesenchymal cells from AML tumors and matched normal kidneys. Green: AML tumor cells; blue: cells from matched normal kidneys; red: tumor-associated fibroblasts (TAF). g. Expression of *MDK*, *GPNMB*, and *NEAT1* in AML tumor cells compared to normal kidney mesenchymal cells and TAFs. The y axis represents the normalized gene expression value. ****p<0.001 (Wilcoxon test). h. Hallmark pathways enriched in AML cells (green) and in matched normal mesenchymal cells (blue). x-axis shows pathway enrichment score. Top five enriched pathways are shown. i. Expression and regulon activity of *SIRT6* and *HDAC2* in tumor and normal mesenchymal cells using same UMAP coordinates of F. Regulon activities of these two transcription factors were calculated based on the expression of their target genes. Note: TAF cells colored in grey were not analyzed for regulon activity. j. RNA *in situ* hybridization (ISH) assessment of *MDK* in AML tumor and in adjacent normal kidney. RNA *in situ* hybridization assessments were performed on the same samples subjected to single cell analysis. An H&E image of the same sample is also provided.

Data integration identified tumor cells and all other expected major cell types in the immune and stromal compartments of AML tumors and matched normal tissues (Fig. 1b-c). Cell types were annotated first by unbiased cross-referencing to two databases of pure cell types, using SingleR^23^, with normalized data. This was followed by manual annotation with cell type specific marker genes to refine cell type identification (Supplementary Fig. 2a). AML and LAM cells were identified using a panel of five well-established marker genes known from prior work to be highly expressed in both AML and LAM^24^ (*CTSK*^25^, *PMEL*^26^, *VEGFD*^27, 28^, *MITF*^29^, and *MLANA*^30^) (Supplementary Fig. 2b). Graph-based clustering was performed on the mesenchymal cell population using Seurat, resulting in eight clusters (Supplementary Fig. 2c). Cells expressing at least two of the five marker genes at or above median expression across all mesenchymal cells with non-zero values were identified as AML/LAM cells. We observed that nearly all cells meeting this criterion were in three clusters (cluster 1, 2 and 6), and therefore, we annotated all cells in these three clusters as tumor cells. The number of AML cells (6%) and LAM cells (0.66%) was low. Importantly, no cells with this expression pattern were observed in the normal kidney specimens, strongly suggesting that this method of tumor cell identification was specific, although it may have undercounted the tumor cell fraction in both AML and LAM. Cells from each patient sample contributed to each cluster, suggesting an absence of major batch effects (Supplementary Fig. 2d-e). Normal kidney contained 49% epithelial cells in contrast to 1.1% epithelial cells in AML, as expected (Fig. 1d). Many immune populations were enriched in tumors compared to matched normal, including macrophages (18.3% vs 2.7%), dendritic cells (4% vs 0.7%), and T cells (32.6 % vs 14.1%). We also identified proliferating T cells and proliferating macrophages in AML (Supplementary Fig. 1a).

The major cell types identified in LAM lung included immune cells (T cells, NK cells, B cells, macrophages and monocytes), mesenchymal cells, epithelial cells and endothelial cells (lymphatic and blood) (Fig. 1e). Proliferating macrophages were also identified in LAM (Supplementary Fig. 2f). In contrast to the AML, no proliferating T lymphocytes were identified in the LAM specimens.

### Global mapping of pathways and genetic regulatory networks in AML cells

Re-clustering of mesenchymal population showed separate clusters of cells from normal kidneys and AML tumors (Fig. 1f). The trimodal cluster consists of cells derived solely from AML tumors. Besides AML cells (as described above), the cluster also contains tumor associated fibroblasts (TAF) with no expression of tumor marker genes but high expression of known TAF marker genes: Tumor-Derived Adhesion Factor (*IGFBP7*)^31^, Fibroblast-Specific Protein-1(*FSP1/S100A4*)^32^, Platelet-Derived Growth Factor Receptor Beta (*PDGFRB*)^32^, Secreted Protein Acidic And Rich in Cysteine (*SPARC*), and SPARC-Like Protein 1 (*SPARCL1*) (Fig. 1f, Supplementary Fig. 3a). TAF have been shown to promote tumor proliferation in many human cancers^33^.

Differential gene expression analysis by Seurat^34, 35^ identified 160 genes uniquely upregulated in tumor cells compared with TAF and normal kidney, including genes previously reported (e.g., *GPNMB*^8^, *SQSTM1*/p62^36^, *MMP2*^37^, *PTGDS*^38^) and novel genes involved in tumor metastasis (e.g. *MMP11*, *MDK*, *DCN*, *PDPN*) (Fig. 1g, Supplementary Fig. 3b, Supplementary data 2). Four long non-coding RNAs (lncRNAs) (*MALAT1*, *NEAT1*, *TUG1*, *MEG3*) were upregulated in both tumor cells and tumor-associated fibroblasts compared to matched normal mesenchymal cells (Fig. 1g and Supplementary Fig. 3c), suggesting remodeling of fibroblasts by AML cells.

To identify pathways differentially regulated in AML cells vs. TAF and normal kidney, we used Gene Set Variation Analysis (GSVA)^39^, a non-parametric, unsupervised method for estimating variation of gene set enrichment. Hallmark gene set analysis (containing 50 gene sets) identified genes involved in cholesterol homeostasis as the most upregulated pathway in AML cells, consistent with prior work^40, 41^, while the second most upregulated pathway was mTORC1 signaling, a well-known biochemical effect of TSC2 loss in AMLs and LAM (Fig. 1h). ROS, glycolysis, and adipogenesis pathways were also enriched in AML, consistent with prior work^36, 42–44^.

To investigate transcriptional networks driving the expression characteristics of AML, we used Single-Cell Regulatory Network Inference and Clustering (SCENIC)^45^. This regulon analysis revealed that more regulons were upregulated in AML cells rather than downregulated. Known TSC-associated transcription factors^9, 46–48^were re-identified, such as *MITF* and *TFE3*, for which both expression and regulon activity were much higher in tumor cells compared to normal kidney mesenchymal cells (Supplementary Fig. 3d). Similarly, *SREBF1*/*SREBF2* and *PPARG*, known master regulators of lipid and cholesterol metabolism downstream of mTORC1^40, 41, 49^, had both high expression and high regulon activities in AML cells (Supplementary Fig. 3d). This analysis also identified novel transcription factors and regulons associated with AML, including several involved in epigenetic regulation, e.g. *HDAC2*, *SIRT6*, *FOXN3, MEF2A* (Fig. 1i, Supplementary Fig. 3d, Supplementary data 3-4).

Specific genes of interest include *MDK* (newly identified here as highly expressed in AML) and *GPNMB* (a known marker of AML^50^), both of which are increased in AML cells in both the scRNA-Seq dataset (Fig. 1g) and the bulk RNA-seq dataset of tumor samples^8^ (Supplementary Fig3. e). Using RNA *in situ* hybridization, *MDK* expression was detected in AML tumors but not in adjacent normal kidney (Fig. 1j), consistent with the single cell data. *MDK* is a direct target of the transcription factor *SP1*^51^, and regulon analysis showed enriched *SP1* expression and activity in the cells with high *MDK* expression (Supplementary Fig. 3f). AML cells with high *MDK* expression showed higher expression of *HIF1A* (Supplementary Fig. 3g), which binds to a hypoxia responsive element in the *MDK* promoter^52^. MDK is an heparin-binding growth factor^53^ that promotes cell growth and angiogenesis^54, 55^.

### AML tumor cells exhibit two major states: stem-like and inflammatory

UMAP re-clustering of the 6,596 AML cells revealed four clusters (Fig. 2a-b). Differential expression analysis revealed high expression of synthetic smooth muscle genes^56^ in cluster 1 and high expression of contractile smooth muscle genes^56^ in cluster 2 (Supplementary Fig. 4a). Clusters 0 and 3 appeared to represent intermediate or transitional cell states between clusters 1 and 2, with a gradient expression of synthetic and contractile smooth muscle marker genes (Supplementary Fig. 4a). Custer 1 showed relatively high expression of the mesoderm-specific transcription factor 21 (*TCF21*), a master regulator of phenotypic modulation of smooth muscle cells^56^ (Fig. 2b, Supplementary Fig. 4b). In disease conditions, phenotypic modulation transforms smooth muscle cells from a differentiated contractile state into a dedifferentiated synthetic state. We noticed that several genes (*SOX4*, *TCF4*) (Fig. 2b, Supplementary Fig. 4c), known to be stem cell markers, were upregulated in cluster 1, and therefore calculated the “stemness score” using a curated list of 50 tumor stemness marker genes^57^. Cluster 1 showed the highest stemness scores which declined in a gradient leading to cluster 2 (Fig. 2c), as well as high activity of signaling pathways involved in stemness including Notch, Hedgehog, and WNT pathways (Supplementary Fig. 4d). Cluster 2 was enriched in immune pathways (Supplementary Fig. 4d), and showed high expression of inflammatory genes including *CCL3*, *CCL4* and *IL1B* (Fig. 2b, Supplementary Fig. 4e). Based on these features, we defined cluster 1 as a stem-like state (SLS) and cluster 2 as an inflammatory state (IS). Differential expression analyses of cluster 1 (SLS) versus cluster 2 (IS) identified 231 differentially expressed genes at fold change >2 (Supplementary Fig. 2f-g). Metabolic kinetic models using generalized mass action (GMA) equations have been used to simulate and predict biological processes^58, 59^. We previously showed that kinetic models of metabolic pathway systems can be used to interpret transcriptomic profiles measured during disease for cellular metabolism modeling^60^. Purine related metabolism is linked to the mTORC1 pathway^61–63^, and high levels of purine nucleotides are required to maintain cancer stemness^64^, while external hypoxanthine supplementation promotes tumor stemness^64^. Therefore, we generated pseudo-bulk RNA-seq data from single cell transcriptomes to infer cellular purine metabolism in both SLS and IS populations as well as normal mesenchymal cells obtained from matched normal kidney in this study. We found that metabolism of guanine/guanosine in the purine pathway was elevated in both tumor cell states compared to normal controls (Fig. 2d). In contrast, hypoxanthine and inosine metabolism were elevated specifically in the SLS population, suggesting that metabolic mechanisms may contribute to the high stemness features seen in the population.

**Fig. 2.**
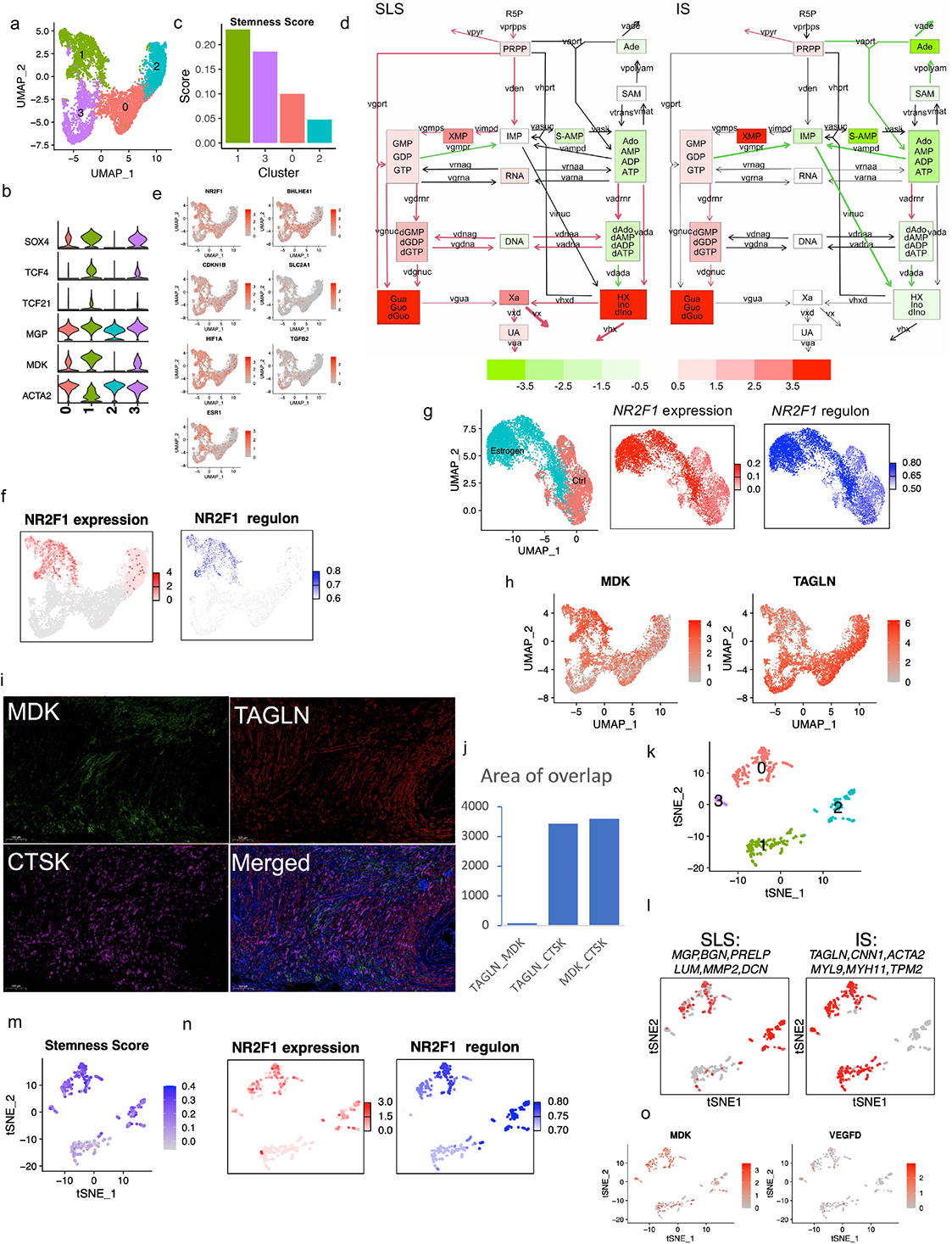
Heterogeneous cellular states in AML and LAM. a. UMAP plot of AML cells only showing two distinct clusters (cluster 1 and cluster 2) and two transitional clusters (0 and 3). b. Violin plots of highly expressed genes of each cluster. The y axis represents the normalized gene expression value. c. Stemness score calculated using 50 tumor stem cell marker genes for each cluster (see method). d. Inferred purine metabolism flux in SLS and IS populations relative to matched normal mesenchymal cells using pseudo-bulk RNA-seq generated from single cell transcriptomes. Relative levels of hypoxanthine (HX) and inosine/deoxyinosine (Ino/dlno) are upregulated in SLS. e. Feature plots of expression of dormancy marker genes in the tumor cell population. f. Analysis of *NR2F1* expression and regulon activity in AML cells. Left panel: expression of *NR2F1* in cluster 1 and 2; right panel: *NR2F1* regulon activity based on 41 downstream target genes. Note: only cluster 1 and cluster 2 were compared for regulon activity. Other clusters are colored in grey. g. *NR2F1* expression and regulon activity in LAM patient-derived TSC2-deficient cells 621-101 cells with and without estrogen treatment. Left panel: t-SNE plot of cells from estradiol treated group (red) and cells from control group (blue); middle panel: expression of *NR2F1*; right panel: *NR2F1* regulon activity. h. Expression of *MDK* and *TAGLN* in AML cell clusters. i. Triple staining for MDK, TAGLN, and CTSK. j. Quantification of co-staining of MDK, TAGLN and CTSK shows little co-localization of MDK and TAGLN (first bar), while both TAGLN and MDK co-localize with the tumor marker gene CTSK (second and third bars). The y axis represents area of overlap (arbitrary unit). k. Re-clustering of the LAM cells from 5 LAM lungs, revealing four clusters. l. Average expression of SLS (left) and IS (right) marker genes in the LAM clusters. An SLS population (cluster 2 in K), IS population (cluster 1 in K), and an intermediate state (Cluster 0 and 3 in K) were identified. m. SLS population (Cluster 2) and intermediate state LAM cells (Cluster 0 and 3) show high stemness scores. n. Expression and regulon analysis of *NR2F1* in LAM cells. Left panel: *NR2F1* expression; right panel: *NR2F1* regulon activity corresponding to the degree of regulation of 23 downstream target genes. o. Feature plot showing expression of *MDK* and *VEGFD* in LAM cells.

The SLS (cluster 1) also showed higher expression of genes associated with TGF-beta signaling and the hypoxia pathway (two main triggers of tumor cell dormancy)^14, 65^, as well as upregulation of the unfolded protein response, which is often seen in dormant tumor cells^14^ (Supplementary Fig.4d). It has been increasingly recognized that a hypoxic microenvironment, as well as stress induced during metastasis, trigger a dormant state in which tumor cells become resistant to drug treatment and stress^66^. Further analysis of a panel of dormancy marker genes revealed high expression in the SLS population (cluster 1), including the transcription factor *NR2F1* (Fig. 2e). NR2F1 serves as a critical node in the induction and maintenance of tumor stem cell dormancy by integrating epigenetic programs of quiescence and survival^14, 67^. Regulon analysis confirmed that *NR2F1* regulon activity (pathway activity of 41 genes regulated by *NR2F1*) was upregulated in the SLS (cluster 1) (Fig. 2f).

Other dormancy marker genes also showed high expression in SLS, including DEC2 (*BHLHE41*), Hypoxia Inducible Factor 1 Subunit Alpha (*HIF1A*), and estrogen receptor alpha (*ESR1*) (Fig. 2e). Estrogen receptor alpha was shown to be required by breast cancer cells to enter NR2F1-dependent dormancy^14^. Hormonal signaling is of particular interest in TSC, since 1) LAM affects almost exclusively women, 2) LAM and AML cells express ER alpha, and 3) estrogen impacts the survival, metastasis, and metabolism of TSC2-deficient cells in models of LAM^68^. To investigate whether ER alpha contributes to dormancy in TSC-deficient settings, as suggested by the scRNA-Seq data, we used TSC2-deficient 621-101 cells^69^, which were derived from a LAM patient’s angiomyolipoma. The cells were treated with 100nM estradiol or vehicle control for 24 hours and subjected to scRNA-Seq. All of the major dormancy genes were upregulated in the estradiol treated group compared to the control group (Supplementary Fig. 2h). The related gene Estrogen Related Receptor Alpha (*ESRRA*) was also elevated by estradiol treatment. Regulon analysis further showed that estradiol treatment increased *NR2F1* expression and regulon activity (Fig. 2g).

The identification of SLS and IS populations was validated in tumor specimens by co-staining with antibodies to SLS and IS markers (*MDK* and *TAGLN* respectively, Fig. 2h), and Cathepsin K (AML/LAM marker gene^25^). MDK positivity was observed primarily in one population, while TAGLN positivity was observed primarily in a separate population (Fig. 2i). As expected, CTSK stained both populations. Quantification revealed little co-localization of MDK and TAGLN, versus extensive co-staining of MDK with CTSK or TAGLN with CTSK (Fig. 2j), supporting the existence of two distinct populations of AML cells, MDK+ and TAGLN+. Staining of MDK, a marker of SLS, for each AML sample is shown in Supplementary Fig. 2i.

### Cell populations occur in LAM that are similar to the two types observed in AML

In the sporadic form of LAM, angiomyolipoma are common, and genetic studies have shown that the AML and LAM cells arise from a common precursor cell^70^. To determine whether the two cell states identified in AML are present in pulmonary LAM, we analyzed 57,186 cells from five LAM lungs using the same marker gene set and method as used for AML. A total of 375 LAM cells were identified (Fig. 1e). Considering the LAM cells alone, UMAP clustering revealed four clusters (Fig. 2k). Similar to AML, one cluster expressed SLS/synthetic smooth muscle marker genes (cluster 2) and another cluster expressed IS/contractile smooth muscle marker genes (cluster 1) (Fig. 2l). An intermediate state with expression of all these genes was also identified (cluster 0). The SLS population and the intermediate state showed higher stemness score (Fig. 2m) and genes associated with dormancy were upregulated in the SLS population and intermediate state (Supplementary Fig. 5a), similar to the SLS cluster in AML. In addition, like the SLS AML cells, the SLS cluster of LAM cells had upregulation of *NF2F1* expression and regulon activity (Fig. 2n). Interestingly, the expression of *VEGFD*, a validated LAM biomarker^71^, was much lower than *MDK* (a potent angiogenic and lymphangiogenic growth factor^55, 72^) in LAM cells (Fig. 2o), suggesting a potential role of MDK in LAM-associated lymphangiogenesis. Thus, we measured MDK serum levels in women with LAM and healthy controls and found that MDK levels were 3.7-fold higher in LAM patients (n = 20) compared to healthy controls (n = 19) (p=0.04, Supplementary Fig. 5b).

### The stem-like population of AML cells may contribute to rapamycin resistance

Rapalog therapy for AML and LAM leads to sustained but incomplete responses, with regrowth of AML and ongoing loss of lung function in LAM when treatment is stopped^6, 7^. These partial responses suggest possible drug tolerance in a subset of AML/LAM tumor cells. Our observation of elevated stemness and dormancy in a subset of tumor cells, typical features of drug-tolerant tumor persister cells^73^, led us to directly examine rapamycin tolerance in AML cells. We developed a primary culture from one of the AML tumors profiled in this study (AML1162 with TSC2 mutation allele frequency of 41%). After one week in culture, these cells were treated with either DMSO (control) or rapamycin for 24 hours followed by scRNA-Seq profiling. A total of 2,066 cells and 4,083 cells were analyzed in the control and treatment group, respectively, after filtering out low quality cells (Fig. 3a). Merging these two sets of cells, UMAP clustering identified seven clusters (Fig. 3b). Marker genes identified in each cluster are provided in Supplementary data 5. Using the same expression criteria described above, a total of 2004 candidate AML cells were identified, accounting for 33% of all cells (Fig. 3c).

**Fig. 3.**
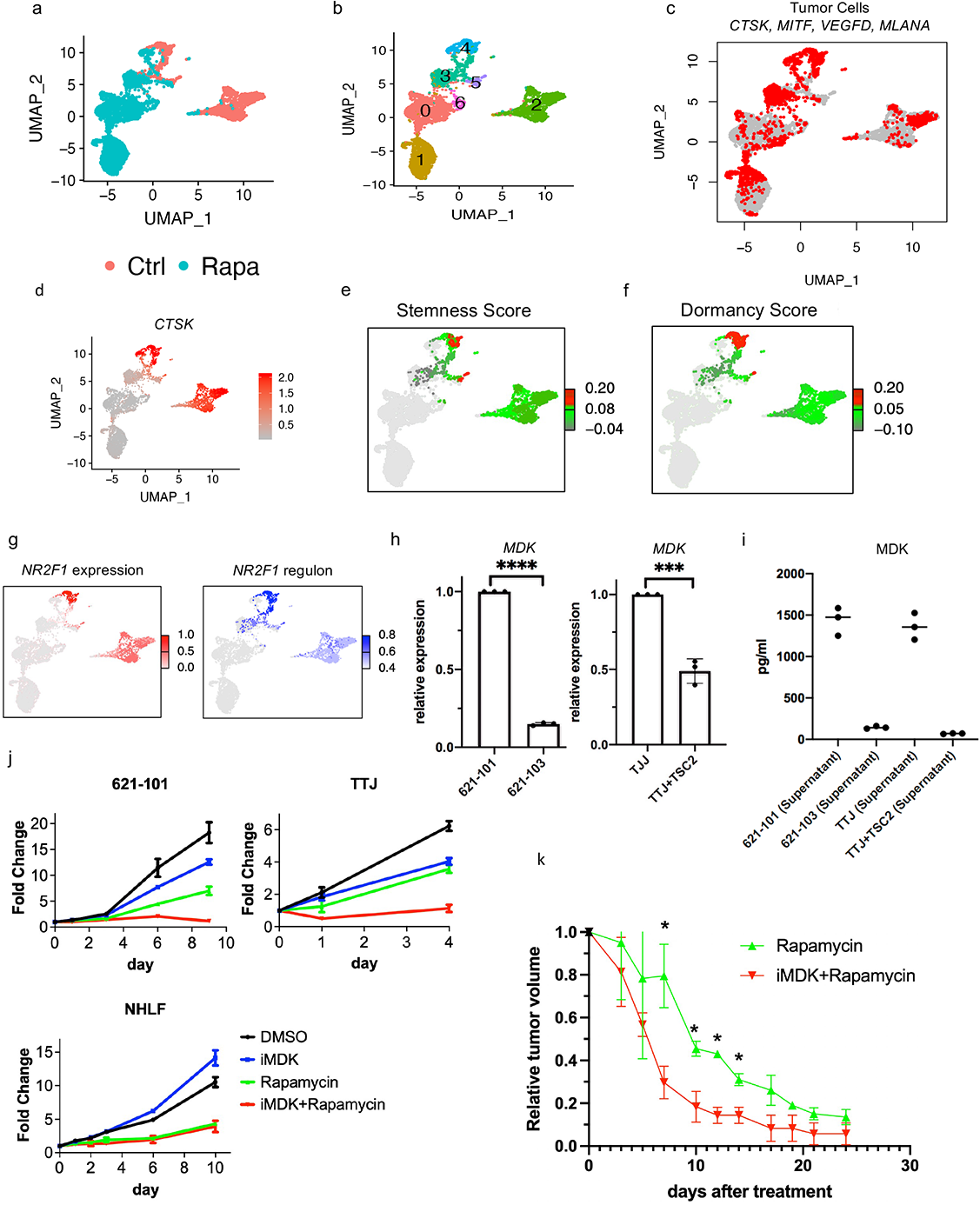
The stem-like population of AML cells may contribute to rapamycin resistance. a. UMAP plot of primary cultures derived from AML tumor colored by treatment. Cells were treated with DMSO as control (red) or 20nM rapamycin (cyan) for 24 hours before scRNA-Seq. b. UMAP plot of the AML-derived primary cell culture, colored by clusters. c. Expression of 5 AML markers in the primary AML culture before and after rapamycin treatment (as in A). d. CTSK expression in cells before and after rapamycin (as in A). e. Cluster 4 in the DMSO control group showed high stemness score, calculated using a panel of 50 cancer stem cell marker genes (see method). Note that stemness score was only calculated in DMSO control group, and the cells in rapamycin treatment group are colored in grey. f. Cluster 4 in the DMSO control group showed high dormancy score, calculated using known dormancy marker genes (see method). Note that dormancy score was only calculated in DMSO control group, and the cells in rapamycin treatment group are colored in grey. g. Expression (left) and regulon activity (right) of *NR2F1* in control group. Note: only cluster 2 and cluster 4 were comparatively analyzed for regulon activity. The color bars indicate expression level and regulon activity only for these two clusters; other cells are colored in grey. h. Relative expresion of *MDK* in TSC2-deficient cell lines compared to TSC2-addback cells. Left panel: patient-derived TSC2-deficent 621-101 cells compared to TSC2-addback 621-103 cells; right panel: mouse kidney derived TSC2-deficent TTJ cells compared to TSC2-addback cells. i. MDK protein level in the cell culture supernatants measured by ELISA. j. Proliferation measured by crystal violet assay. Treatments were: DMSO, 1µM iMDK, 20nM rapamycin, and combination of iMDK and rapamycin for days indicated. 621-101 and TTJ are TSC-deficient. NHLF: normal human lung fibroblasts. k. Tumor size reduction relative to pre-treatment tumor volume in rapmayin treatment and combined iMDK and rapamycin treatment groups. * p<0.05 (t-test). Relative tumor size after treatment for all treatment groups can be found in Supplementary Fig. 6d: TTJ xenograft mice (n=6 per group) were treated 3 times/wk with DMSO, iMDK (9mg/kg), rapamycin (3mg/kg), or combined iMDK (9mg/kg) and rapamycin (3mg/kg). Averaged tumor size was reduced to <20% of pre-treatment volume after 4 treatments in the combination treatment group, in contrast to 8 treatments in the rapmaycin treatment group.

Rapamycin had a striking effect on overall transcriptomes, and most clusters were composed nearly entirely of either treated or untreated cells. Strikingly, we identified a small cluster (cluster 4) that contained AML cells from both control and rapamycin treatment groups, suggesting that it contained cells that are resistant to rapamycin, or at least cells in which transcription was not changed by rapamycin treatment. In this cluster, the expression of many AML tumor marker genes was unaffected by rapamycin, in contrast to other clusters where rapamycin suppressed expression of these tumor genes (Supplementary Fig. 6a), including *CTSK* (Fig. 3d).

Further analysis of cluster 4 showed high expression of tumor marker genes (Supplementary Fig. 6b, only DMSO control group is shown in the UMAP), with a strikingly similar expression pattern to that of the SLS population of AML tumors. For instance, we have identified elevated expression of *SOX4*, *PTGDS*, *MMP2* among other marker genes in AML tumors (Supplementary Fig. 4f), suggesting that cluster 4 corresponds to the SLS state of AML cells. In addition, cells in the cluster 4 showed a high stemness score (Fig. 3e), and high dormancy score (calculated by expression of known dormancy marker genes^14^ including the dormancy inducer *NR2F1* and hormonal regulator *ESR1*) (Fig. 3f and Supplementary Fig. 6c). Consistent with these results, *NR2F1* regulon activity was high in this cluster (Fig. 3g).

Levels of the dormancy inducer *NR2F1* and the hormonal regulator *ESR1* (ERa) were unchanged by rapamycin (Supplementary Fig. 6a), suggesting that dormancy may be associated with treatment resistance. Expression of *SQSTM1* (p62) and *SOD2*, which help to maintain cellular ROS homeostasis in TSC^36^, were unaffected by rapamycin in cluster 4 (Supplementary Fig. 6a), suggesting that redox homeostasis maintenance may be involved in treatment tolerance. These data suggest that the SLS state is resistant to rapamycin treatment, which is consistent with the notion that acquired stemness and dormancy render tumor cells resistant to chemical therapeutics^74–76^.

MDK is reported to mediate drug resistance in other tumors^77^, and we observed high expression of *MDK* in the SLS population (Fig. 2h). To determine whether MDK is involved in rapamycin tolerance and whether MDK is regulated by TSC pathway, we used two cellular models of TSC and found that expression of *MDK* was upregulated in TSC2-deficient AML patient-derived 621-101 cells compared to TSC2-reexpressing 621-103 cells, as well as in mouse kidney derived TSC2-deficient TTJ cells^78^ compared to TSC2-addback TTJ+TSC2 cells (Fig. 3h). Because MDK is a secreted cytokine, we further assessed MDK protein levels in the cell culture medium by ELISA. MDK levels were significantly higher in both the patient-derived and mouse-derived TSC2-deficient cell lines compared with TSC2-addback controls (Fig. 3h).

Next, to assess the importance of MDK expression on rapamycin resistance *in vitro* and *in vivo*, we used an MDK inhibitor (iMDK) that specifically inhibits MDK but not other growth factors such as VEGF or pleiotrophin (PTN) (homologous to MDK)^79^ and was shown to potently inhibit MDK and thus enhance PD-1 therapy in melanoma mouse models^80^. TSC2-deficient cells (621-101, TTJ) and normal human fibroblasts (NHLF) were treated with DMSO, rapamycin (20nM), iMDK (1µM), or a combination of rapamycin (20nM) and iMDK (1µM). Treatment with iMDK alone had minimal effects in all 3 cell lines. However, when combined with rapamycin, iMDK had a synergistic effect on the two TSC2-null cell lines (Fig. 3j). We defined synergy as the combined effect of two drugs is greater than the sum of each drug’s individual activity^81, 82^. In normal fibroblasts (NHLF), rapamycin had a dramatic growth inhibitory effect, which was not significantly changed by the addition of iMDK. To determine whether iMDK sensitizes tumors to rapamycin treatment *in vivo*, we generated subcutaneous tumors using the TSC2-deficient TTJ cells in immune-deficient athymic nude mice. Combination treatment with iMDK and rapamycin led to a more rapid onset of tumor response, and a lower tumor burden, compared with rapamycin alone, while iMDK alone had no apparent effect (Fig. 3k, Supplementary Fig. 6d). Many cancers have relatively high MDK expression in comparison to matched normal tissues, including bladder cancer (Supplementary Fig. 6e). We found that three bladder cancer cell lines were also sensitized by iMDK to rapamycin treatment (Supplementary Fig. 6f). These data may provide a rationale for combination therapy targeting MDK and mTORC1 in TSC and other selected tumors.

### Remodeling of endothelial cells by heterogeneous tumor cell states

We next investigated the potential differential effects of these two cell states, SLS and IS, on the tumor microenvironment. As seen in Fig. 1, both blood and lymphatic endothelial cells were enriched in AML compared with adjacent normal kidney, suggesting ongoing angiogenesis and lymphangiogenesis. Strikingly, the distribution of IS and SLS was not uniform among our six AML samples, with two AMLs consisting mainly of IS (> 70%), and four mainly SLS (>80%) (Fig. 4a). The SLS-dominant tumors had a much greater content of endothelial cells, with an average of 24.9% fenestrated endothelial cells and 1% lymphatic endothelial cells, in contrast to the IS-dominant tumors with average 1.4% fenestrated endothelial cells and 0.6% lymphatic endothelial cells (Fig. 4b). To validate this, immunohistochemistry (IHC) staining for the endothelial marker CD31 was performed on each AML and the percentage of endothelial cells was calculated by digital analysis, revealing a strong correlation with the percentage predicted by scRNA-Seq (Fig. 4c) and a higher percentage of endothelial cells in SLS-dominant tumors (Fig. 4d, Supplementary Fig. 7a). This dramatic difference in the endothelial composition in SLS-dominant vs. IS-dominant tumors suggests that the endothelial cells are responding to specific cues arising from the predominant cell type within the tumor. Thus, we investigated all genes that were over-expressed in SLS compared to IS and again identified *MDK* as the top differentially expressed angiogenic gene (by fold change). VEGF-D (a pro-lymphangiogenic factor) is thought to drive pulmonary lymphangiogenesis in LAM. Differential expression analysis showed much higher expression of *MDK* than *VEGFD* in general, with higher expression of *MDK* in cluster 1 (SLS) and higher expression of *VEGFD* in cluster 3 (IS) (Fig. 4e). Interestingly, *VEGFA,* another well-recognized pro-angiogenic factor was only expressed in a small number of SLS cells (Supplementary Fig. 7b). Taken together, these data suggest that high expression of pro-angiogenic MDK in SLS tumors may account for the enriched endothelial cells in this subtype of AML.

**Fig. 4.**
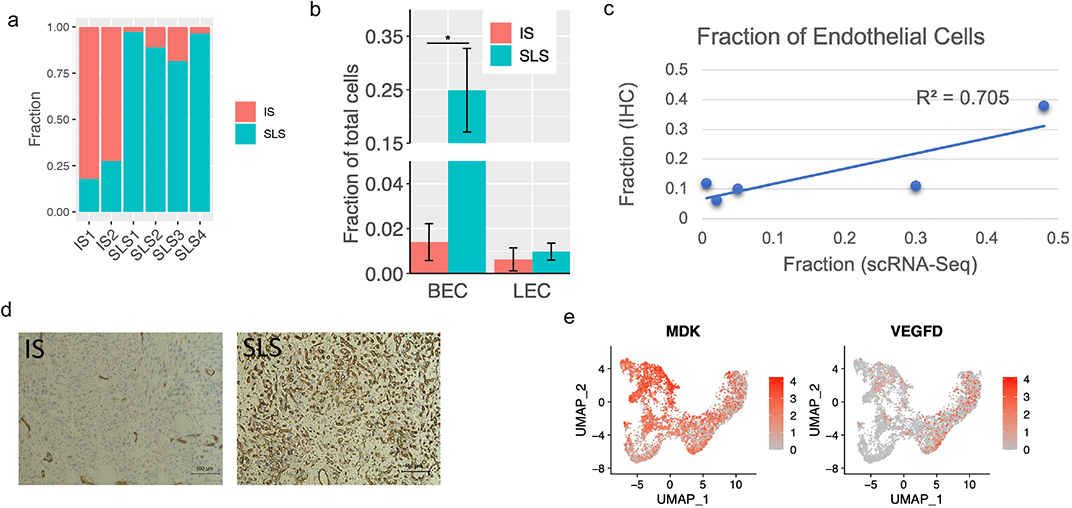
Endothelial cell remodeling in SLS-dominant tumors. a. Percentage of SLS and IS cells in the six AML tumors profiled. b. Quantification of fractional representation of blood endothelial cells (BEC) and lymphatic endothelial cells (LEC) in SLS-dominant (n=4) and IS-dominant tumors (n=2). Standard errors are shown for each group. *p < 0.05, two-sided Wilcoxon test. c. Comparison of percentage of blood endothelial cells identified by single cell profiling and CD31 IHC for five AML tumors. d. Representative IHC staining of endothelial cells with anti-CD31 in IS-dominant tumor and in SLS-dominant tumor. e. Expression of *MDK* and *VEGFD* in AML cell populations. The MDK figure is also shown in Fig. 2h, and is repeated here for ease of comparison.

Further differential and pathway analysis revealed remodeling of endothelial cells in AML, including high expression of C-C Motif Chemokine Ligand 21 (*CCL21*), *TBX1* and *NRP2* specifically observed in tumor LECs (Supplementary Fig. 7c-g). Regulon analysis further revealed that transcription factors *NR2F1* and *NR2F2* may underlie these transcriptional programs (Supplementary Fig. g-h).

### T cell dysfunction and suppressed clonal expansion in SLS-dominant tumors revealed by integrative analysis of scRNA-Seq and scTCR-Seq

T cell infiltration and exhaustion have been observed in human TSC tumors, and a clear benefit of immunotherapy was observed in mouse models^17, 18^. To determine whether T cells are influenced by tumor cell states in AML, we focused on the four AMLs with paired normal kidneys profiled, two of which were SLS-dominant and two of which were IS-dominant. Pathway activity analysis of tumor-derived T cells compared to that from paired normal kidneys revealed upregulation of inflammatory responses, including the type I and type II interferon pathways (Supplementary Fig. 8a). Cell proliferation pathways (E2F targets, MYC targets, Mitotic signaling) were also consistently upregulated in tumor-derived T lymphocytes, in line with the observed general expansion of T cells (Fig. 1d). A population of proliferating CD8+ cells (CD8 T-prolif) was present specifically in the tumors (Supplementary Fig. 8b-c) and not in normal kidney, suggesting an expansion of tumor antigen-reactive T lymphocytes. T cell expansion in tumors was confirmed by CD3 IHC (Supplementary Fig. 8d). Multiple immune checkpoint markers were expressed in tumor derived T cells (Supplementary Fig. 8e).

The higher fraction of T cells in IS-dominant tumors compared to SLS-dominant tumors suggests more T cell infiltration and/or T cell proliferation in IS-dominant tumors (Supplementary Fig. 8f), consistent with previous reports that stem-like states in tumors are associated with immunoresistance^83^. Re-clustering of tumor-derived CD8+ T cells (down-sampled to have equal number of cells from SLS or IS samples) revealed three major clusters: memory/naïve T cells (CD8 Tm/naïve), effector T cells (CD8 Teff) and proliferating T cells (CD8 T-prolif), as well as subclusters within each major cluster, with different expression of immune checkpoint genes or cytotoxic effector genes (Fig. 5a-b). We calculated an exhaustion score for each cell based on relative expression of known checkpoint genes, including T Cell Immunoreceptor With Ig And ITIM Domains (*TIGIT*), Lymphocyte Activating 3 (*LAG3*), B- and T-Lymphocyte Attenuator (*BTLA*) and Killer Cell Lectin Like Receptor G1 (*KLRG1*); and a cytotoxic score based on relative expression of cytotoxic effectors, including Granzyme B (*GZMB*), Interferon Gamma (*IFNG*) and Tumor Necrosis Factor (*TNF*). CD8+ T cells derived from SLS-dominant tumors showed much lower cytotoxic scores compared to those derived from IS-dominant tumors, and a lower percentage of cytotoxic cells (defined as expressing at least one cytotoxic effector genes) within each subpopulation (Fig. 5c). In addition, SLS-dominant tumor derived cells exhibited higher exhaustion scores (Fig. 5c). Despite a roughly equal frequency of exhausted cells in each subpopulation (Fig. 5c), the fraction of exhausted CD8+ Teff cells in SLS-dominant tumors was significantly higher than that in IS-dominant tumors, and IS-dominant tumors showed a higher frequency of both cytotoxic CD8+ Teff and CD8+ Tm/Naïve populations (Fig. 5d). Similar analysis of tumor-derived CD4+ T cells revealed six subtypes of CD4+ T cells (Fig. 5e-f). While memory CD4+ T cells and CD40LG-high population derived from IS-dominant tumors showed a higher cytotoxicity score (Fig. 5g), no significant difference in cell frequency in any subtype was observed between SLS-dominant and IS-dominant tumors (Fig. 5h).

**Fig. 5.**
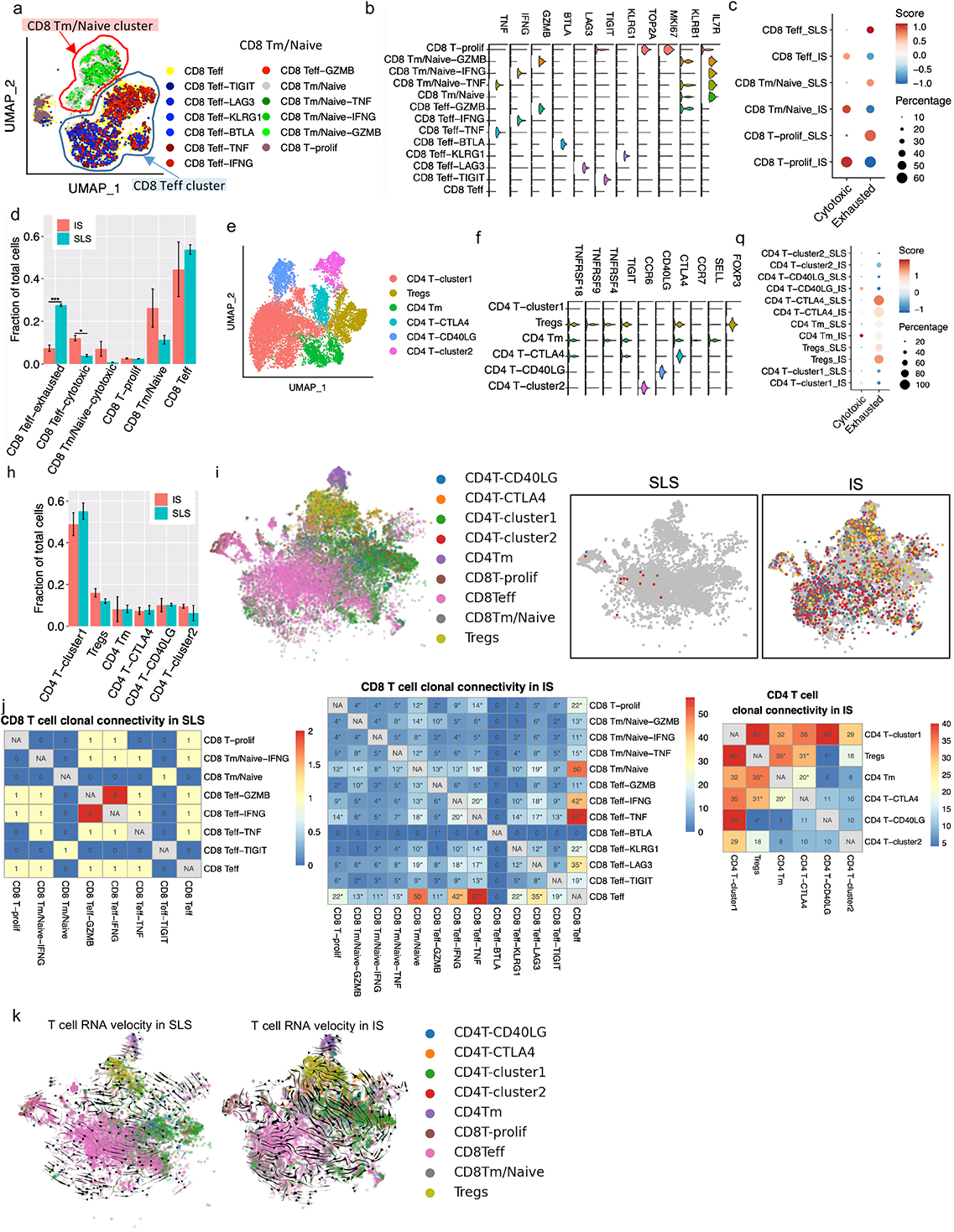
T cell dysfunction and suppressed T cell clonal expansion in SLS-dominant tumors. a. UMAP plot of CD8+ T cells obtained from four AML tumors (downsampled to have equal number of cells from SLS or IS dominant tumors). Phenotypic clusters are represented in distinct colors. CD8 Teff: effector CD8+ T cells; CD8 Tm/Naïve: memory/naïve CD8+ T cells; CD8 T-prolif: proliferating CD8+ T cells. b. Violin plot of representative marker genes of each cluster of CD8+ T cells defined in (A). The y-axis represents the normalized gene expression values. c. Module score of T cell exhaustion or cytotoxicity across major CD8+ T cell population in SLS or IS dominant tumors. Exhaustion module score was calculated based on relative expression of checkpoint genes *TIGIT*, *LAG3*, *BTLA* and *KLRG1.* Cytotoxicity module score was calculated based on relative expression of cytotoxic effector genes *GZMB*, *IFNG* and *TNF*. Module scores were scaled with red color representing higher score. Percentage of cytotoxic or exhausted cells in each population is represented by the circle size. d. Quantification of fractional presentation of clusters of CD8+ T cells across two subtypes of tumors. Standard errors are shown for each group. *p < 0.05, **p<0.01, ***p<0.001, t-test. e. UMAP of clusters of CD4+ T cells obtained from four AML tumors (downsampled to have equal number of cells from SLS or IS dominant tumors). f. Violin plot of representative marker genes of each cluster of CD4+ T cells. The y-axis represents the normalized gene expression value. g. Module score of T cell exhaustion or cytotoxicity across major CD4+ T cell population in SLS or IS dominant tumors calculated as C. h. Quantification of fractional presentation of clusters of CD4+ T cells across the two subtypes of tumors. Standard errors are shown for each group. No significance difference was identified. i. Representative shared T cell clonotypes identified in IS-dominant tumor and in SLS-dominant tumor. Each clonotype is represented by a different color. Major cell groups are display on left panel. j. Shared TCR clonotypes in CD8+ T cells and CD4+ T cells, after normalizing to total cell numbers. Number of shared clonotypes between each pair of subtypes were displayed. *p < 0.05, two-sided Fisher’s exact test. No shared clonotypes were identified in CD4+ T cells in SLS-dominant tumor. k. RNA velocity of T cell population calculated based on ratio of unspliced and spliced transcripts in each cell. Velocity vectors represented by arrows indicate potential differentiation paths.

Tumor activated lymphocytes undergo clonal expansion, and expanded T cells from the same clone have the same TCR sequence (clonotypes), which enables tracking of differentiation trajectories. We examined sharing of expanded TCR clonotypes across all sub-populations of CD8+ and CD4+ T cells within individual samples after cell number normalization, which revealed 229 clonotypes shared among CD8+ T cell subtypes and 319 clonotypes shared among CD4+ T cell subtypes in IS-dominant tumors but only 5 clonotypes shared among CD8+ T subtypes in SLS-dominant tumors (Fig. 5i). While SLS tumors showed quite limited clonotype sharing among subtypes in CD8+ T population and no clonotype sharing in CD4+ T population, IS tumors exhibited extensive clonotype sharing among subtypes in both CD8+ T and CD4+T populations (Fig. 5j): 75% of expanded TCRs in the CD8+ Teff subtype were shared with the CD8+ Tm/Naïve subtype in IS tumors, revealing a dynamic connection between these two CD8+ T cell states. In IS tumors, the majority of proliferating CD8+ T cells shared clonotypes with CD8+ Teff population, which may imply a tumor antigen-reactive T cell proliferation (Fig. 5j). Proliferating T cells shared a high number of clonotypes with two cytotoxic Teff populations (CD8 Teff-TNF and CD8 Teff-IFNG). In addition, extensive clonal sharing was observed between CD8 Teff and two cytotoxic Teff populations (CD8 Teff-TNF and CD8 Teff-IFNG), suggesting an active and dynamic differentiation trajectory toward functional T cells. These observations suggest that the high frequency of cytotoxic CD8+ T cells observed in IS tumors is at least partially due to a dynamic differentiation of presumably tumor-recognizing effector cells. As expected based on previous work^84^, CD4+ T cells showed less clonal expansion in general compared to CD8+ T population. CD4+ T cells clonal sharing was only detected in IS-dominant tumors. The Tregs cluster shared TCRs with CD4 T-cluster1, CD4 Tm and CD4 T-CTLA4 clusters (Fig. 5j), suggesting a complex dynamic differentiation of Tregs in tumors.

To infer dynamic differentiation among subtype T cells, we calculated splicing-based RNA velocity using single cell transcriptome data^85^. Consistent with the substantial clonal connectivity observed in IS-dominant tumors, this analysis supported a differentiation trajectory from CD8 effector T cells to proliferating T cells and from multiple CD4+ T subpopulations to Tregs (Fig. 5k). In contrast, SLS-dominant tumors showed limited differentiation potential among subtypes.

Given the striking difference in T cell modulation in SLS versus IS dominant tumors, we next explored whether SLS tumor cells express higher levels of immune checkpoint genes to inhibit T cells. We analyzed TIGIT ligands (*PVR*, *NECTIN2*), BTLA ligand (*TNFRSF14*), LAG3 ligand (*HLA-DRA*, *FGL1*), *KLRG1* ligand (*CDH1*, *CDH2*), and PD-1 ligands (*CD274*, *PDCD1LG2*). Surprisingly, all of these ligands showed low expression in both groups of tumor cells (Supplementary Fig. 8g). The low expression levels and lack of significant differences of these immune checkpoint ligands between SLS versus IS tumors suggest other mechanisms in the differential modulation of T cell function in these tumor cell states.

### Delineating the suppressive immune microenvironment in TSC

Immunosuppressive myeloid cells, such as tumor-associated macrophages (TAMs), are considered major barriers to cancer immunotherapy^86^, due to their potent suppressive function and high abundance in the tumor microenvironment^87^. As noted above, enrichment of macrophages represented the most striking immune infiltration in AML (Fig. 1b,1d). This enrichment of macrophages in the AML was confirmed by CD68 IHC (Fig. 6a). These AML-derived macrophages showed higher expression of the immune checkpoint genes T cell immunoglobulin and mucin domain-containing protein 3 (TIM3) encoded by *HAVCR2*, and V-domain immunoglobulin suppressor of T cell activation (VISTA) encoded by *VSIR*, in comparison to macrophages derived from matched normal kidneys (Fig. 6b). The expression of other immune checkpoint genes is provided in Supplementary Fig. 9a. Expression of VISTA and TIM3 on tumor infiltrating macrophages is associated with T cell dysfunction in the tumor microenvironment^88, 89^.

**Fig. 6.**
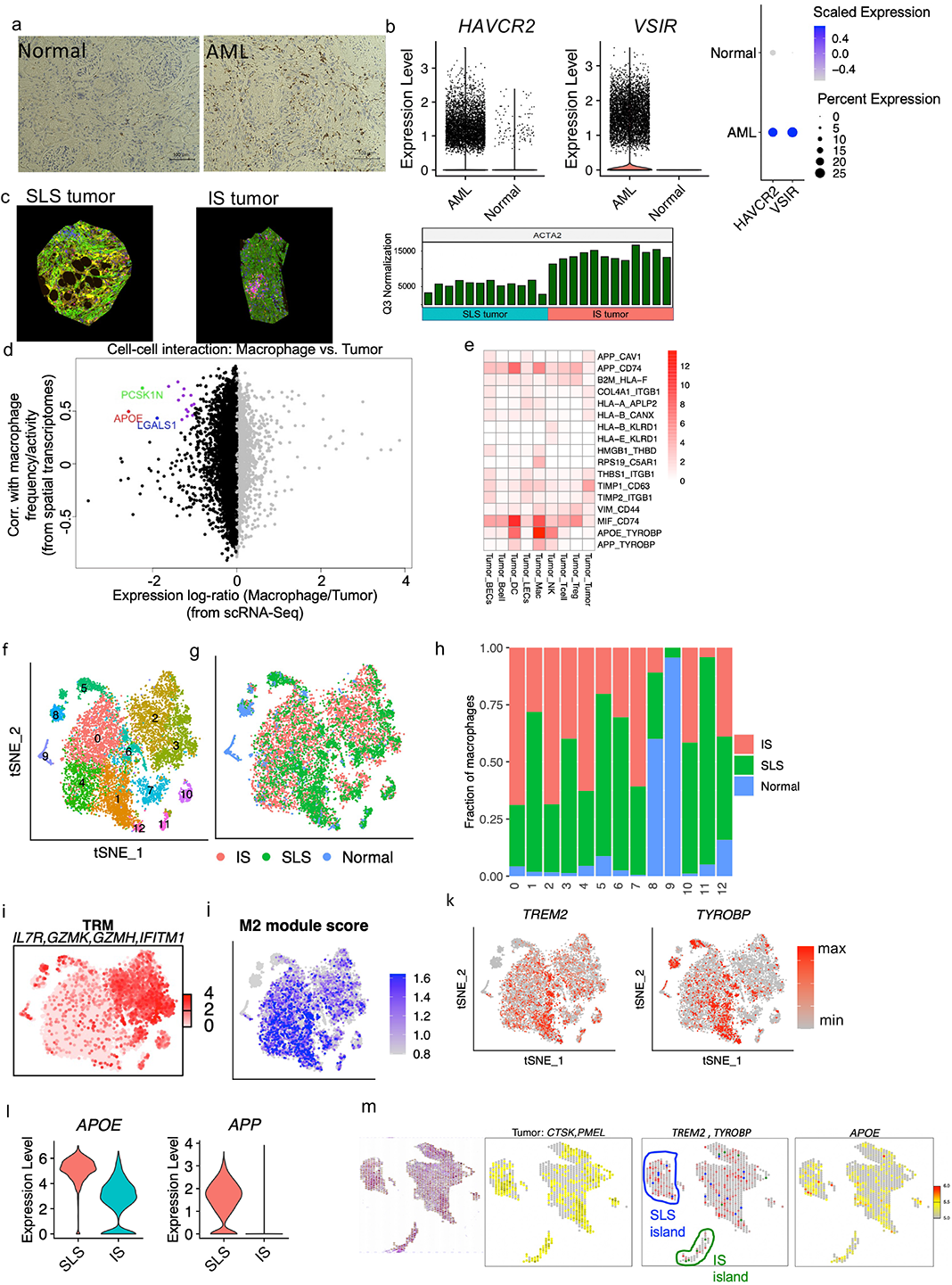
The suppressive immune environment is shaped by tumor cell states. a. CD68 IHC staining on a representative AML tumor and matched normal kidney. b. Higher expression of TIM3 (*HAVCR2*) and VISTA (*VSIR*) in macrophages obtained from tumors compared to macrophages obtained from matched normal kidneys. Left panel: violin plot showing expression of *HAVCR2* and *VSIR*; right panel: dot plot showing scaled expression and the percentage of cells expressing these genes. c. Nanostring digital spatial profiling of one SLS-dominant and one IS-dominant tumor. Left panel: a representative ROI (Region of Interests) from SLS-dominant tumor; middle panel: a representative ROI from IS-dominant tumor; right panel: expression of ACTA2 across all ROIs after Q3 normalization (From left to right columns are 12 SLS-dominant tumor ROIs and 12 IS-dominant tumor ROIs). d. Inferred interactions between tumor cells and macrophages calculated by integrative analysis of spatial transcriptomics of the representative SLS-dominant tumor (12 ROIs) and scRNA-seq. x-axis displays relative expression of genes in single cell data. Only genes that are expressed in both single cell data and spatial transcriptomics data are shown. Left side are genes relatively highly expressed in tumor cells; right side are genes relatively highly expressed in macrophages. Y-axis displays Pearson Correlation Coefficient (PCC) of gene expression with macrophage frequency in spatial transcriptomics data. Genes with log-ratio less than −1.5 and correlation coefficient higher than 0.4 are colored. *APOE*: PCC=0.49, p=0.1 (correlation test). e. Interactions between tumor cells and other cell types calculated as the product of the average ligand expression and average receptor expression (only interactions with a score greater than 1 across any cell type pair are displayed). Each column shows a pair of cell types, and each row shows the ligand-receptor pair. The color indicates interaction score. Column label: cell type expressing the ligand and cell type expressing the receptor are separated by “_”. Row label: ligand and receptor are separated by “_”. f. tSNE plot of macrophages colored by cluster (downsampled to have equal number of cells from SLS or IS tumors). g. tSNE plot of macrophages colored by sample type of origin. h. Fraction of macrophages obtained from subtypes of tumors or from matched normal across clusters. i. Average expression of *IL7R*, *GZMK*, *GZMH*, and *IFITM1*. j. M2 module score calculated by relative expression of *CD163*, *MRC1*, *VEGFA*, *GPNMB* and *TREM2*. k. Feature plot showing expression of *TREM2* and *TYROBP* (DAP12). l. Violin plot showing expression of *APOE* and *APP* in tumor cells (SLS vs. IS). m. Spatial transcriptomic profiling of an independent AML tumor using 10x Visium platform. Panels from left to right: 1) H&E stained tissue, 2) averaged expression of *CTSK* and *PMEL*; spots with expression of CTSK and PMEL higher than the median of all spots were annotated as tumor (yellow), 3) SLS spots (blue) and IS spots (green) were identified by marker gene expression; averaged expression of *TREM2* and *TYROBP* are displayed in red, 4) expression of *APOE*.

Tumor cells may influence other cells in the microenvironment by direct ligand-receptor interactions or indirect cell-to-cell communications in which tumor cells produce a signal (such as paracrine effectors) to recruit or exclude immune cells and alter their behavior^86, 90^. Therefore, we further analyzed one SLS tumor and one IS tumor using Nanostring digital spatial profiler (DSP) to query spatial tumor-microenvironment organization, and confirmed higher expression of the IS marker gene *ACTA2* in IS-dominant tumors (Fig. 6c). For each tumor, we selected 12 regions of interest (ROIs) that were enriched with tumor cells (smooth muscle actin positive, green), T cells (CD3 positive, pink), and macrophages (CD68 positive, yellow) for RNA sequencing (Supplementary Fig. 9b). Since single cell data have much higher resolution, we re-defined gene signatures (Supplementary data 6) for each of the major cell types identified in AML, including macrophages, Langerhans cells, dendritic cells, T cells, B cells, lymphatic endothelial cells and blood endothelial cells. We then used these cell type specific gene signatures to deconvolute the cell composition of each selected area and to infer the relative frequency or activity of these cell types in each of the ROIs, and searched for genes primarily expressed by tumor cells that may influence or correlate with the frequency/activity of another cell type. We reasoned that genes expressed primarily by tumor cells may influence a different cell type in the tumor microenvironment by an indirect paracrine signal, hence correlation analysis of the expression of genes (primarily expressed in tumor cells) and frequency/activity of another cell type in each ROI was performed to reveal genes mediating cell-to-cell communication, as shown previously^90^. We found a high correlation of *APOE*, *LGALS1* and *PCSK1N*, which were primarily expressed in SLS tumor cells, with macrophage frequency/activity in SLS-dominant tumors (Fig. 6d, Supplementary data 7). These correlations were not observed in IS-dominant tumors (Supplementary Fig. 9c). Interestingly, all of these genes encode secreted proteins, suggesting a specific paracrine regulatory role of SLS tumor cell secretome on macrophages. Consistent with this concept, APOE and LGALS1 were previously shown to promote M2 polarization of macrophage/microglia in mouse models^91, 92^. We next analyzed a published bulk RNA-Seq dataset of ten AML tumors^8^, which again revealed a high correlation between *APOE* and macrophage population frequency (Supplementary Fig. 9d). Since bulk RNA-Seq data are confounded by tumor purity and tumor heterogeneity, the robust identification of this correlation strongly supports the existence of a tumor-macrophage regulatory axis.

To search for putative macrophage receptors for these tumor ligands and to profile the full spectrum of ligand-receptor mediated direct tumor-microenvironment interactions, we next performed ligand-receptor interaction analysis using a validated algorithm previously described^86^ and a list of over 2,500 curated pairs of ligand-receptors to infer putative tumor-microenvironment interaction based on ligand expression in one cell type and corresponding receptor expression in another cell type. This revealed tumor-macrophage interactions via *APOE*-*TYROBP* (DAP12) as the strongest interaction among tumor-microenvironment interactions (Fig. 6e). TYROBP and TREM2 form a receptor complex on macrophages which has been extensively studied in the context of neurodegenerative diseases, where the complex mediates signaling and cell activation following binding to its ligands including APOE or β-amyloid (a cleavage product of the amyloid-beta precursor protein APP)^93–95^. Interestingly, *APP* also showed strong interaction with *TYROBP*. Recent studies have shown that TREM2+/TYROBP+ tumor-associated macrophages (TAMs) suppress T cell function and proliferation in various tumors and that targeting this TAM population can modulate immunosuppressive TAMs and restore T cell function^96, 97^.

To compare macrophages derived from SLS and IS tumors, we down sampled to 5,000 cells from each tumor type and included all macrophages derived from matched normal kidneys for downstream analysis. Re-clustering identified 12 clusters (Fig. 6f). Most clusters were primarily derived from tumors except two small clusters (cluster 8 and cluster 9) that were mainly derived from normal kidneys (Fig. 6g-h). Two main types of macrophages (tissue resident macrophages and TAMs) were identified in the tumor derived macrophage population. Cluster 2 and cluster 3 were annotated as tissue resident macrophages (TRM) based on high expression of IL7R^98^ and inflammatory genes (Fig. 6i and Supplementary Fig. 9e). The TAMs in AML are mainly composed of 4 clusters (cluster 0, cluster 1, cluster 4, and cluster 6) characterized by a high M2 module score, which was calculated by the relative expression of alternatively activated macrophage marker genes, including CD163^99^, MRC1^99^, VEGFA^100^ and TREM2^96^ (Fig. 6h and Supplementary Fig. 9f). Surprisingly, the organization of TAMs showed a striking difference between SLS and IS tumors: cluster 1 and cluster 6 were mainly composed of cells derived from SLS tumors, whereas cluster 0 and cluster 4 were mainly composed of cells derived from IS tumors (Fig. 6i). Cells from cluster 1 and cluster 6 showed high expression of *TREM2* and *TYROBP* (Fig. 6k). These data show that there is a higher percentage of *TREM2*+/*TYROBP*+ TAMs derived from SLS tumors.

These observations suggest a regulatory axis from SLS tumor cells to TAMs via an APOE-TREM2/TYROBP interaction, with APOE as a putative ligand for the TREM2/TYROBP complex in tumor TME. Consistent with this hypothesis, *APOE* (and *APP*) showed higher expression in SLS AML cells compared to IS cells (Fig. 6l). We sought to validate this observation using 10x Visium spatial transcriptomic profiling in an independent AML sample. We used *CTSK* and *PMEL* to identify AML cells (Fig. 6m). Spots with averaged expression of *CTSK* and *PMEL* higher than 50% across all spots were annotated as tumor spots(yellow). We then calculated scores for SLS and IS within identified tumor spots using the most robust marker genes *MGP* (for SLS) and *ACTA2* (for IS) (see Methods), and identified islands enriched with SLS (blue) or IS (green). Plotting average expression of *TREM2* and *TYROBP* (red) revealed higher expression in the SLS enriched island compared to IS enriched island (Fig. 6m). APOE also showed higher expression in the SLS enriched island (Fig. 6m).

Our discovery of a striking suppression of CD8+ T cells in SLS-dominant tumors is consistent with the reported role of *TREM2*+/*TYROBOP*+ TAMs in suppressing CD8+ T cell function and proliferation in tumors^96^. The higher T cell clonal expansion and dynamic differentiation in IS-dominant tumors suggest tumor-reactive T cell activation. Taken together, this tumor-specific inhibition of T cell function and T cell proliferation/differentiation in SLS-dominant AML implies a major immunomodulatory role of myeloid cells in TSC. This has particular importance given the extremely low expression of immune checkpoint ligands on the AML tumor cells (Supplementary Fig. 8g).

### Analysis of molecular interactions between tumor and tumor microenvironment provides potential targets for distinct precision therapeutic strategies for SLS and IS tumor

In the immune compartment, we also observed enrichment of B lymphocytes and dendritic cells in AML relative to normal kidney. We detected 1,620 B cells predominately from tumor (Fig. 7a-b). Re-clustering revealed six clusters. Of these, five were particularly tumor enriched. We identified follicular B cells expressing high levels of CD20 (*MS4A1*) and *CXCR5* in both tumor (cluster 5) and adjacent normal kidneys (clusters 1) (Fig. 7b). In contrast, plasma B cells expressing immunoglobulin gamma (*IGHG1*, *CD27*, *CD38*) were exclusively enriched in tumors (Fig. 7c). Pathway analysis identified induced interferon gamma and TGF beta signaling in regulatory B cells, suggesting a regulatory role of Tregs in tumor microenvironment^101^ (Supplementary Fig. 10a). A pattern of reduced activity in tumor-specific plasma B cells, evidenced by a universal downregulation of pathways involved in cell growth (Myc targets, mTOR pathway) and inflammation (interferon alpha/gamma, IL2 and TNF alpha signaling), may suggest reduced function of plasma cells in the tumors.

**Fig. 7.**
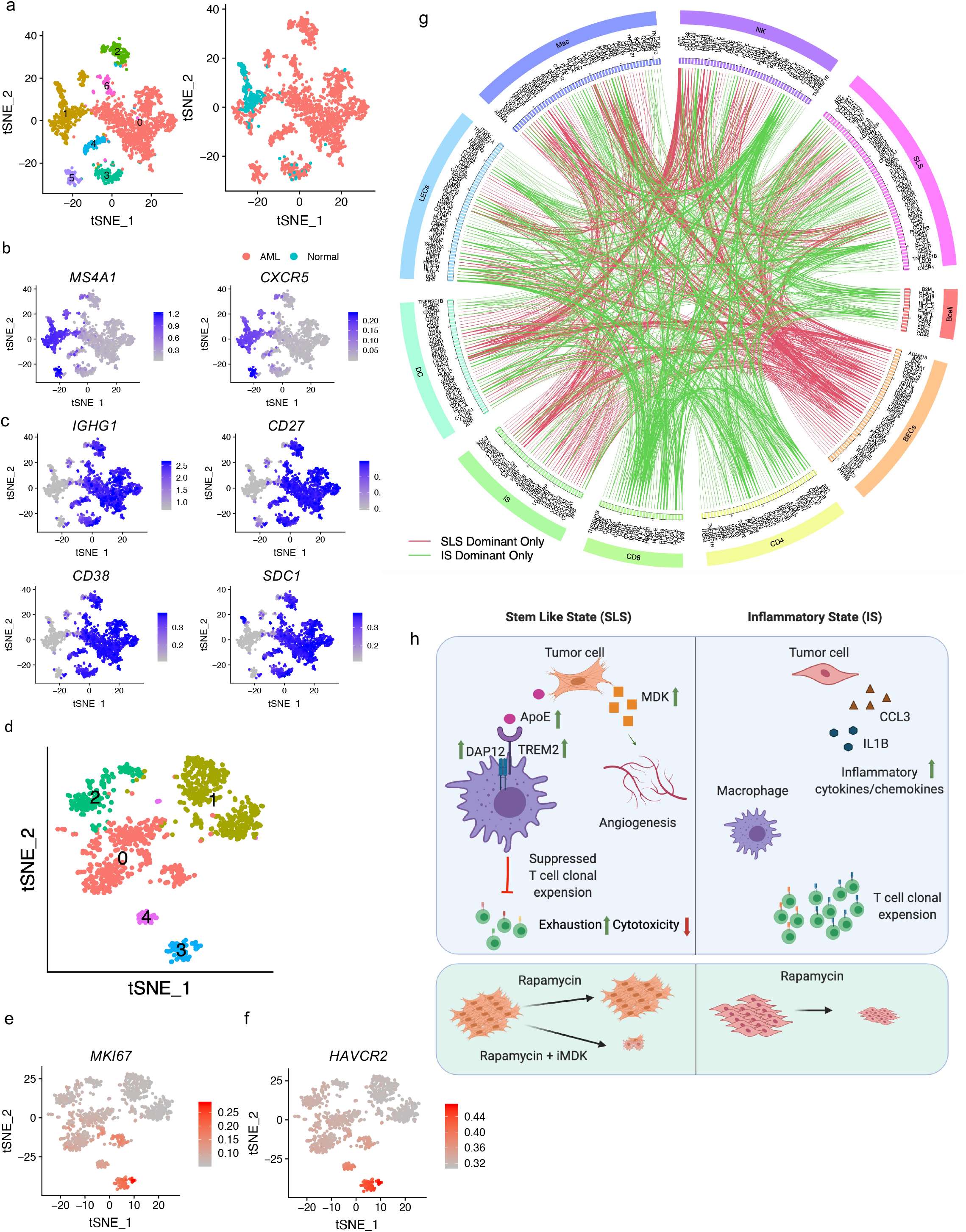
Molecular interactions between tumor and immune compartment inferred by ligand-receptor co-expression. a. tSNE plot of 1,620 B cells colored by cluster (left) or the origin (right). b. Feature plot showing expression of follicular B cell marker genes *MS4A1* and *CXCR5*. c. Feature plot showing expression of plasma B cell marker genes. d. tSNE plot of dendritic cells from AML tumors which are colored by cluster. e. High expression of MKI67 in proliferating dendritic cells. f. High expression of TIM3 (*HAVCR2*) in proliferating dendritic cells. g. Circos-plot showing ligand-receptor pairs identified across pairs of cell types (cutoff value for interaction is 1). Red lines indicate pairs only identified in SLS-dominant tumors; green lines indicate pairs only identified in IS-dominant tumors. h. Schematic showing the main discoveries from this study: identification of two cell states (SLS and IS), their differential cellular ecosystem with active crosstalk between tumor cells and microenvironment, and association with rapamycin resistance and immune modulation. In SLS tumor cells, upregulated APOE may modulate tumor-associated macrophages toward an immune suppressive state by directly binding to TREM2/TYROBP receptor complex, leading to T cell dysfunction and diminished T cell clonal expansion; upregulated MDK expression may induce angiogenesis and drive persistence in response to mTORC1 inhibition. MDK is identified as a novel therapeutic target combining with rapamycin for persisting SLS tumor. In contrast, IS tumors with upregulated inflammatory pathways exhibit higher T cell cytotoxicity/proliferation and sensitivity to rapamycin treatment.

We detected 839 cross-presenting dendritic cells expressing *CLEC9A* and *XCR1* (Supplementary Fig. 10b) exclusively in tumors. Re-clustering identified a small cluster of proliferating cells (cluster 3) (Fig. 7d-e). This cluster showed higher activity of Myc targets, E2M targets and mTORC1 signaling (Supplementary Fig. 10c). *HAVCR2* (TIM-3) has been reported to be an important regulatory factor of dendritic cells in anti-tumor immunity^102, 103^. The proliferating cluster exhibited high *HAVCR2* expression (Fig. 7f), suggesting a pro-tumoral function of proliferating dendritic cells.

Although AML has an extremely low mutational burden^19^, the overall enrichment of plasma B cells and cross-presenting dendritic cells in tumors may suggest tumor-specific antigen presentation in tumor microenvironment that may include all the genes/proteins highly expressed in AML, including CTSK and MDK.

Tumor-microenvironment interactions play crucial roles in tumor development^104^. To assess the comprehensive crosstalk between tumor and tumor microenvironment, we quantified potential cell-cell interactions among all cell types in the tumor microenvironment as described above. We observed numerous interactions in SLS-dominant tumors, including B2M-HLA-F, HLA-B-CANX, and MIF-CD74 in B and T cells (Fig. 7g and Supplementary Fig. 10d-e), similar to what has been reported for melanoma^86^, which is of interest because as described above, AML express many melanoma marker genes including MITF, PMEL and MLANA^46, 47^. Interestingly, we identified more tumor-TAF interactions in SLS-dominant tumors (21 pairs) compared to IS-dominant tumors (4 pairs) by ligand-receptor analysis (Supplementary data 8). We also observed extensive interactions of KLRD1 and HLA family members between NK cells and other cell types (Supplementary Fig. 10f), and interactions between tumor cells and other cell types related to extracellular matrix remodeling. Tumor cell secreted extracellular matrix molecule such as collagen (COL4A1) can bind to adhesion receptors broadly expressed on many cell types, such as integrin receptor ITGB1 (Supplementary Fig. 10g). We observed expression by tumor cells of thrombospondin (THBS1) and tissue inhibitors of metalloproteinases (TIMP1 and TIMP2), secreted factors involved in extracellular matrix remodeling (Supplementary Fig. 10g).

Differential analysis of landscape of ligand-receptors interactions in SLS-dominant versus IS-dominant tumors revealed different tumor-microenvironment crosstalk in these two tumor cell states. For example, more interactions between tumor and blood endothelial cells were found in SLS-dominant tumors, consistent with enriched endothelial cells in SLS-dominant tumors (Fig. 7g). The depletion of interactions of CD8 and CD4 T cells with other cell types in SLS-dominant tumors may underlie the molecular mechanisms for the observed suppressed T cell clonal expansion (Fig. 7g).

## Discussion

mTORC1 is estimated to be hyperactive in at least half of all human malignancies and plays a central role in tumorigenesis^105–107^. Our work provides a comprehensive atlas of tumor cells and the tumor microenvironment in mTORC1 hyperactive AML and LAM. Our analysis highlights a complex cellular ecosystem with active crosstalk between AML cells and the tumor microenvironment and distinct AML/LAM cell states associated with rapamycin resistance and immune modulation (Fig. 7h). In addition to confirming known genes and pathways contributing to TSC pathogenesis, we highlight previously unrecognized pathways that likely contribute to tumor progression, and pinpoint novel targets for the future of immunotherapy in TSC. Our study represents an important step toward understanding intra-tumoral expression heterogeneity in mesenchymal tumors, a far less studied tumor type than epithelial tumors.

Among our key findings is the identification of a conserved drug resistant tumor cell state characterized by stemness and dormancy seen in both AML and LAM. Rapamycin and its analogs induce a cytostatic effect in TSC treatment, resulting in some shrinkage and then stabilized tumor volume. Here, we reveal two distinct cell states (SLS and IS) in the tumor cell population, and identify underlying transcription factors that may drive the development of these different cell states in response to the tumor microenvironment, characterized by distinct expression of tumor stem cell and dormancy programs or inflammatory programs. Immunofluorescent staining confirmed the existence of these cell states, as predicted by single cell transcriptomic profiling. SLS cells with stemness and dormancy properties contribute to rapamycin tolerance as shown by our *in vitro* treatment analyses. Inhibition of MDK, a gene highly expressed in SLS cells, enhanced rapamycin’s therapeutic effect both *in vitro* and *in vivo*, suggesting that MDK may at least partially account for the molecular mechanism of rapamycin tolerance in TSC, in line with role of MDK in drug resistance observed in other cancers^77, 108^. Thus, intra-tumoral heterogeneity, which is believed to underlie therapy resistance in many malignant tumors, also occurs in mTORC1-hyperactive AML and LAM, and combinatorial targeting mTORC1 and factors such as MDK that contribute to this heterogeneity may enhance the efficacy of mTORC1 inhibition.

SLS-dominant tumors were enriched for both blood endothelial cells and lymphatic endothelial cells when compared to IS-dominant tumors, indicating differential induction of vascular remodeling of endothelial cells. We validated this enrichment of endothelial cells by IHC. Lymphatic vascularization is a hallmark of both AML and LAM, AML cells can metastasize to regional lymph nodes, and it has been proposed that LAM cells metastasize to the lungs from a distant unknown site-of-origin^24, 109^ VEGFD is thought to promote lymphangiogenesis and lymphatic metastasis^24^. Serum VEGFD levels are elevated in about two-thirds of LAM patients, serving as an important diagnostic biomarker^110^. Whether other growth factors may contribute to lymphangiogenesis in LAM, including the one-third of LAM patients without elevated VEGF-D, is a critical unanswered question. We identified MDK as a secreted factor that may promote lymphangiogenesis and angiogenesis in SLS-dominant tumors, and found that MDK is elevated in the serum of LAM patients, suggesting that it may be a critical mechanistic link to lymphangiogenesis in LAM as well as a candidate therapeutic target.

Compared to matched normal kidneys, a higher percentage of T cells was observed in AML tumors, and proliferating T cells were solely observed in tumors, indicating tumor-induced T cell activation and expansion. This concept is supported by increased expression of genes associated with inflammation in tumor-associated T cells revealed by comparative pathway analysis. This T cell infiltration in tumors was validated by IHC and supports the conclusion of a prior study of T cells in AML^17^. Evidence of T cell exhaustion was present in the effector T cell population, consistent with T cell exhaustion previously reported in human AML and LAM and in mouse models^17, 18^, which may curtail the proliferation and cytotoxicity of tumor-recognizing T cells^111^. Intriguingly, CD8+ T cells derived from SLS-dominant exhibited much higher exhaustion and lower cytotoxicity compared to those from IS-dominant tumors. Integrative analysis of paired scRNA-Seq and scTCR-Seq revealed that clonal expansion and T cell velocity were almost completely suppressed in SLS-dominant tumors.

We observed striking macrophage infiltration in these renal AML, validated by IHC and consistent with previous observations in hepatic AML^21^, emphasizing a possible role of the innate immune system in TSC. M2 polarization of TAMs is implicated in tumor promotion and immune suppression^112^. A subset of M2-like TAMs was observed in AML, characterized by high expression of M2 marker genes. Interestingly, it seems that macrophage alternative polarization in AML tumors is shaped by different tumor cell states. Specifically, SLS-dominant tumors were enriched with M2-like macrophages with high expression of *TREM2* and *TYROBP*, a receptor complex on macrophages recently shown to suppress T cell function in tumor microenvironment^96, 97^. Because TREM2+/TYROBP+ tumor-infiltrating macrophages inhibit T cell proliferation in animal models of sarcoma, colorectal cancer, and mammary tumor^96, 97^, it is possible that these suppressive macrophages are responsible for the observed T cell dysfunction and almost complete suppression of T cell clonal expansion and differentiation observed in SLS-dominant tumors. Integrative analysis of spatial transcriptomic profiling and single cell analysis identified a connection between *APOE* (primarily expressed by tumor cells) and macrophage population frequency, which was robustly recapitulated by a further integrative analysis of bulk RNA-Seq and single cell analysis. Genome-wide ligand-receptor analysis revealed *APOE*-*TYROBP* as the strongest tumor-microenvironment interaction, suggesting a regulatory axis from tumor cells to suppressive TAMs. The TREM2/TYROBP complex acts as a receptor for amyloid-beta protein 42, a cleavage product of the amyloid-beta precursor protein APP^93^ and APOE^113^. Consistently, both *APP* and *APOE* showed higher expression in SLS AML cells compared to IS type. Since expression of known immune checkpoint ligands was extremely low on tumor cells in both SLS-and IS-dominant tumors, T cell function and proliferation/differentiation may be inhibited indirectly by the SLS tumor cells via induced suppressive TAMs. While tumor mutation burden has been associated with response to immune checkpoint therapy in multiple cancer types, it is not a perfect marker of response, and suppressive myeloid cells have gained attention as a critical determinant of therapeutic resistance in multiple cancer types^114^. Our work suggests that in TSC tumors, which are known to have an extremely low mutational burden^19^, suppressive myeloid cells may drive immune suppression, and blocking tumor-myeloid cell crosstalk may provide enhance immune regulation of these tumors.

The relatively small sample size in this study, despite the large number of tumor cells analyzed (6,596 cells), is an important limitation for our findings. Additional analysis with larger numbers of samples for these rare diseases will help to validate these findings. However, we note that the very similar findings in AML and LAM provide some degree of cross-validation, including robust identification of the same two distinct cell states: stem cell like and inflammatory. Interestingly, a recent study found a positive correlation between MDK expression and poor prognosis in melanoma patients and revealed that MDK-educated melanoma secretome promotes immunosuppressive macrophages leading to T cell dysfunction^80^. Given the striking similarity between TSC tumors and melanoma, such as high expression of melanocyte genes *CTSK* and *MITF* and similar tumor-microenvironment crosstalk revealed by this study, the regulatory axis from SLS tumor cells to T cell dysfunction via *TREM2*+/*TYROBP*+ M2-like macrophages identified in this study might be a conserved mechanism across tumor types.

In addition to macrophages, B cells were enriched in tumors compared to matched normal kidneys. Detailed analysis revealed a general downregulation of cell growth and inflammation pathways in tumor infiltrating B cells, which suggests exhaustion of those B cells. In addition, cross-presenting dendritic cells were enriched in tumors compared to matched normal kidneys. Cross-presenting dendritic cells play an important role in the tumor microenvironment by priming T cells to target tumor cells. A group of proliferating dendritic cells were identified, presumably induced by tumor recognition, that showed high expression of immune checkpoint TIM3, likely contributing to the suppression of T cell priming. Finally, mapping molecular interactions between tumor and tumor microenvironment in TSC highlights conserved tumor-microenvironment interactions and potential therapeutic targets. For example, ligand-receptor mediated tumor-microenvironment interactions recapitulate many interactions observed in melanoma. Strikingly, indirect cell-cell interaction mapping revealed that interactions between T cells and other cell types were primarily enriched in IS tumors, whereas interactions of endothelial cells were primarily enriched in SLS tumors. Therefore, in the future, it may be possible to personalize therapeutic design based on these molecular interactions.

In summary, our work provides an atlas of neoplastic, stromal, and immune cells with important and novel insights into TSC biology. The link between the drug-resistant stem cell like state and suppressed T cell dynamics via tumor-educated macrophages revealed by this work has important translational implications. The insights revealed here may have broad relevance for understanding the molecular mechanisms underlying other mTORC1 hyperactive tumors.

## ACKNOWLEDGEMENTS

This work was supported by a Career Development Award from The LAM Foundation (LAM0142C01-20) to Y. T; a grant from the DOD Tuberous Sclerosis Complex Research Program (W81XWH1910152-TS180029) to Y.T; and U01HL131022 (NIH) to E.P.H. We also thank the Rothberg Family and the Engles Family for their support of this work. We thank Dr. Chin-Lee Wu at Massachusetts General Hospital for providing AML specimens. We thank Dr. Krinio Giannikou for downloading and sharing the bulk RNA-seq data published by MacKeigan group^8^. We thank Dana-Farber/Harvard Cancer Center in Boston, MA, for the use of the Specialized Histopathology Core, which provided histology and immunohistochemistry service. Dana-Farber/Harvard Cancer Center is supported in part by an NCI Cancer Center Support Grant # NIH 5 P30 CA06516.

## AUTHOR CONTRIBUTIONS

Y.T. designed and performed experiments, analyzed data and wrote the manuscript. Y.T., D.J.K and E.P.H conceived the project and revised the manuscript.

## METHODS

### EXPERIMENTAL MODEL AND SUBJECT DETAILS

#### Patient samples

LAM specimens, AML tumor samples and matched normal kidneys were collected under IRB approved by the Brigham and Women’s Hospital (Protocol #2008P002071). All patients provided informed consent. None of these patients received rapalog treatment for six months prior to surgery. AML samples were obtained locally from Massachusetts General Hospital, Brigham and Women’s Hospital/Dana-Farber Cancer Institute or Beth Israel Deaconess Medical Center in Boston. LAM samples were obtained either locally from Brigham and Women’s Hospital or from National Disease Research Interchange (NDRI). All samples were immediately dissociated and subjected to single cell analysis upon receipt. No specific sampling was performed for AML or LAM samples. The entire piece was analyzed for scRNA-seq.

#### Cell Lines

The following cell lines were maintained in our lab: patient-derived TSC2-deficient cell line 621-101, TSC2-addback cell line 621-103, mouse kidney derived TSC2-null cell line TTJ (the gift of Vera Krymskaya) and TSC2-add back cell line TTJ-TSC2. The normal human lung fibroblasts NHLF (CC-2512) was purchased from Lonza Group (Switzerland). All cells were cultured in DMEM supplemented with 10% FBS (Thermofisher Scientific), and were cultured at 37°C in a humidified chamber with 5% CO_2_ during the experiments.

#### Mice

Animal studies were approved by the Brigham and Women’s Hospital Animal Care and Use Committee (IACUC). All husbandry and experiment procedures with mice were conducted in accordance with protocols approved. Mice were provided water and food *ad libitum* and were housed on a standard light (12h) and dark (12h) cycle. Athymic nude mice (Crl:NU(NCr)-*Foxn1^nu^*, Charles River Laboratories, Wilmington, MA) were all seven-week-old female mice at the time of TTJ cell injection for allograft experiments.

### METHOD DETAILS

#### Single cell RNA-Seq

##### Tissue dissociation

Fresh tumor and matched normal samples were dissociated into single cell suspension using human tumor dissociation kit (Miltenyi Biotec) and gentleMACS™ Dissociator (Miltenyi Biotec), according to manufacture’s manual. Red blood cells were removed by Red Blood Cell Lysis Solution kit (Miltenyi Biotec). Cell suspensions were washed with cold PBS. Viability of all samples were confirmed with trypan blue staining (Invitrogen) to be above 70% before loading to 10x Chromium controller.

##### Single cell RNA sequencing

Droplet emulsions were immediately recovered for reverse transcription reaction using Bio-rad thermocycler. Single cell expression libraries were constructed using 10x genomics Chromium 5’ barcoding reagents (v1) following manufacturer’s manual. Quality of amplified cDNA and constructed libraries were confirmed by BioAnalyzer (Agilent, High Sensitivity DNA Kit). Library sequencing was performed by NextSeq 500 (Illumina). QC files are provided in supplementary data 9.

##### Paired single cell TCR sequencing

Aliquots of 2 μl amplified cDNA from the single cell expression library construction workflow were used for TCR library construction according to 10x genomics manufacturer’s manual. Quality of amplified cDNA and constructed libraries were confirmed by BioAnalyzer (Agilent, High Sensitivity DNA Kit). Library sequencing was performed by NextSeq 500 (Illumina). Sequencing depth for V(D)J enriched libraries were at least 5000 read pairs per cell. Standard Illumina sequencing primers were used for both sequencing and index reads following 10x manufacturer’s protocol.

##### Nanostring whole transcriptome digital spatial profiling

Two tumors were profiled using Nanostring digital spatial profiler for whole transcriptome analysis. Resected tumor samples were washed with cold PBS and fixed in 50 ml 10% Formalin for 24 hours before embedding. All antibody staining and whole transcriptome sequencing were performed on freshly cut FFPE slides. Regions of interest (ROIs) were selected based on immunofluorescence staining using Nanostring validated antibodies against α-SMA (green), CD3 (pink) and CD68 (yellow). Each ROI was uniquely indexed then pooled for sequencing. Sequencing data were Q3 normalized by a standard Nanostring pipeline. Q3 (3rd quartile of all selected targets) normalization was used for all targets that are above the limit of quantitation. Q3 normalization uses the top 25% of expressed genes to normalize across ROIs/segments.

##### 10x Visium spatial transcriptomics profiling

Fresh tumor sample and matched normal kidney were immediately OCT embedded within 1 hour after surgery. OCT blocks was cryosectioned at −10 °C and placed on chilled Visium tissue optimization slide (PN: 3000394, 10x Genomics) or spatial gene expression slide (PN: 2000233, 10x Genomics). Slides were kept chilled during sectioning and transportation processes. H&E staining, tissue optimization and gene expression library construction were performed as per manufacturer’s manual. Briefly, tissue permeabilization time was set to 18 minutes for gene expression experiment after time-course optimization experiment following manufacturer’s protocol. Brightfield H&E images were taken using Keyence BZX800 microscope with a 20x objective. Images were stitched by Keyence BZX800 stitching function. Fluorescent images were taken with dsRed2 filter cube from Chroma Technology (ex/em: 545/30, 620/60) using a 10x objective. Libraries were sequenced on illumina NovaSeq 6000 at 300 pM concentration.

##### Multiplex immunofluorescence, RNA *in situ* hybridization and immunohistochemistry

FFPE tissue blocks were freshly cut at thickness of 5 μm. The following primary antibodies were used for multiplex immunofluorescence staining performed by iHisto company: MDK [EP1143Y] (ab52637, Abcam) (FITC labeled, green), TAGLN (ab14106, Abcam) (cy5 labeled, red) and CTSK (PB9856, Boster) (cy3 labeled, pseudo-colored pink for visualization). Nuclei were stained with DAPI. Slide images were scanned at 10x magnification.

RNA *in situ* hybridization (ISH) was performed at Brigham and Women’s Hospital Pathology Core according to ACD user manual using RNAscope® 2.5 LS Probe - Hs-MDK-O1 (586478, ACD, Bio-techne). All RNA *in situ* hybridization experiments were performed on the same samples subjected to single cell analysis.

All immunohistochemistry staining were performed at Brigham and Women’s Hospital Pathology Core with validated antibodies against CD68 (clone PG-M1, M0876, Dako), CD3 (clone F7.2.38, A0452, Dako), CD31(clone JC70A, M0823, Dako).

##### MDK ELISA assay

Human and mouse MDK ELISA assays were performed using Human Midkine ELISA Kit PicoKine™ (EK1235, BOSTER) and Mouse MDK / Midkine (Sandwich ELISA) ELISA Kit (LS-F12048-1, LSBio) respectively following manufacturers’ manuals. Standards were prepared immediately prior to performing the experiment. For human patient samples, frozen serum samples were thawed to room temperature and centrifuged at 15 minutes at 1000 x g immediate before assessment. For cell culture assays, 621-101 cells, 621-103 cells, TTJ cells and TTJ-TSC2 cells were cultured to 90% confluence. Culture supernatants were collected, centrifuged at 500 x g for 5 minutes, and assayed immediately.

##### Quantitative PCR assay

Probes Hs00171064_m1 (human, ThermoFisher), Mm00440280_g1 (mouse, ThermoFisher), and TaqMan™ Universal Master Mix II, with UNG kit (4440038, ThermoFisher) were used for Quantitative PCR assays.

##### Allograft tumors

Seven-week-old female athymic nude mice (Crl:NU(NCr)-*Foxn1^nu^*, Charles River) were subcutaneously injected with 3 million TTJ cells (75µl cells mixed with 75 μl Matrigel, Corning 356237) on the front flank. All treatments started at day 8 after tumor inoculation when average tumor volume reached around 300 mm^3^. Mice were randomized in groups for treatment. Mice were treated 3 times per week for a total of 11 treatments with intraperitoneal injections of DMSO vehicle, the small molecule 3-[2-[(4-Fluorophenyl)methyl]imidazo[2,1-b]thiazol-6-yl]-2H-1-benzopyran-2-one (iMDK; TOCRIS Bio-techne, 9mg/kg), rapamycin (Sirolimus A8167, APExBIO; 3mg/kg), or combined iMDK (9mg/kg) and rapamycin (3mg/kg). Tumor volume was measured immediately before each treatment using a caliper.

##### Rapamycin treatment and scRNA-Seq of tumor-derived primary culture

Resected tumor tissue was dissociated into single cell suspension as described above. Aliquots of 100μl were dispensed into 10cm dishes with fresh DMEM supplemented with 10% FBS. Primary cultures were maintained at 37°C in a humidified chamber with 5% CO_2_ for 2 weeks to allow to reach 80% confluence. Fresh media were changed every 3 days. Primary cultures were treated with rapamycin (20nM) or vehicle for 24 hours before subjecting to droplet scRNA sequencing as described above.

##### Cell line estradiol treatment and scRNA-Seq

Patient-derived TSC2-deficient 621-101 cells were grown in phenol-free DMEM supplemented with 10% charcoal-stripped FBS for 72 hours, then treated with 100nM estradiol or ethanol vehicle for 24 hours, and subjected to single cell RNA sequencing as described above.

##### Combination treatment and proliferation assay of cell lines

621-101 cells, TTJ cells, or normal human fibroblasts NHLF cells were seeded in 12-well plates in DMEM with 10% FBS at 20-30% confluency. Cells were treated with DMSO (control), iMDK (1μM), rapamycin (20nM), or combined iMDK (1μM) and rapamycin (20nM) until the control group reached over 100% confluency. Drugs were refreshed every 2 days to ensure maximum activity. Cell proliferation was assessed using Crystal violet Assay Kit (Cell viability) (ab232855, abcam).

### QUANTIFICATION AND STATISTICAL ANALYSIS

Statistical analyses were performed with R, MATLAB, or GraphPad Prism (GraphPad Software). Statistical parameters are reported at appropriate places in main text, supplemental materials, figures and figure legends, including sample numbers, measures of center, standard deviation, or standard error (mean ± SD or SEM), statistical significance.

#### Single cell RNA sequencing data processing

Cell Ranger pipeline (10x Genomics) was used for reference genome alignment and generating gene-cell counts matrices. Raw sequencing data was aligned to GRCh38 reference genome using Cell Ranger pipeline (10x Genomics) to generate gene counts matrix by cell barcodes. Sequencing depth was on average 30,285 reads/cell. Data normalization and integration were performed using the Seurat R package (v3.0.0)^34^. Cells were filtered from downstream analysis with the criteria of < 200 genes or > 6000 genes detected and > 0.1 fraction of mitochondrial gene. Samples were normalized individually and integrated with the IntegrateData function. The integrated Seurat object was further scaled by regressing out UMI count and fraction of mitochondrial genes. Optimal principal components used for dimensionality reduction was determined empirically for each analysis by the drop off in PC variance. Cell cycle regression was not performed given small proliferating cell clusters identified in this study. Differential gene expression was analyzed using Seurat ‘FindAllMarkers’ or ‘FindMarker’ functions.

#### Cell type annotation

We first used an automatic cell type annotation R package SingleR^23^ to annotate cell types. Briefly, this algorithm computes the spearman correlation between the transcriptome of the test cell and reference data (i.e., bulk RNA-seq of a pure cell type or cell state) to define cell type label. The reference datasets used in this study include Human Primary Cell Atlas (HPCA) and Blueprint-Encode. We then manually refined cell type annotation based on marker genes identified using unsupervised clustering and differential expression analyses^34^. All cell type marker genes used in this study were from literature. Expression of major cell type marker genes are shown in Supplementary Fig. 1.

#### Tumor cell population analysis

Tumor cells were identified as expressing at least two of the five literature reported marker genes^25–30^ above median value across all mesenchymal cells with non-zero values. Clustering and tumor cell state annotation were performed using normalized raw data. Tumor stemness score was calculated using Seurat AddModuleScore function based on relative expression of 50 tumor stem cell marker genes described previously^57^.

#### T cell population analysis

Four AML tumors and paired normal kidneys were analyzed for T cell function, two of which were SLS-dominant and two of which were IS-dominant. T cell population was downsampled to have equal number of cells from SLS or IS samples. We calculated an exhaustion score for each cell based on relative expression of known checkpoint genes, including *TIGIT*, *LAG3*, and *KLRG1*; and a cytotoxic score based on relative expression of cytotoxic effectors, including *GZMB*, *IFNG* and *TNF*. Cells with expression of at least one checkpoint gene or one cytotoxic effector gene were calculated for the scores and were regarded as exhausted or cytotoxic respectively.

#### Single cell T cell receptor and T cell clonotype analysis

Raw FASTQ reads were mapped to human GRCh38 V(D)J reference genome (v3.1.0, 10x Genomics) using Cell Ranger pipeline (10x Genomics). Sequencing depth was on average 20,876 reads/cell. The filtered contig annotation file was used for downstream analysis that contains high-confident contigs. For clonotype analysis, we downsampled to roughly equal number of cells derived from SLS and IS tumors. After normalizing the cell numbers, we detected 4,667 unique clonotypes in two IS-dominant tumors and 220 unique clonotypes in two SLS-dominant tumors. Clonotype size ranged from 1 to 632 cells in IS-dominant tumors and 1 to 23 cells in SLS-dominant tumors. We further defined clonotype expansion as that a clonotype shared by at least three cells within individual sample, and clonotype sharing as that a clonotype detected in any two or more T cell subtypes within individual sample. we detected that 69% of clonotypes were expanded in IS-dominant tumors, and 18% clonotypes were expended in SLS-dominant tumors. In IS-dominant tumors, we identified 229 and 319 shared clonotypes in CD8+ and CD4+ T cells respectively, whereas, in SLS-dominant tumors, we identified 5 and 0 shared clonotypes in CD8+ and CD4+ T cells respectively. One-sided Fisher’s exact test followed by Benjamini-Hochberg correction was used to assess statistical significance of clonotype sharing among T cell subtypes on cluster-by-cluster contingency tables.

#### RNA velocity analysis

RNA velocity was calculated using scVelo (v0.2.2, python package)^85^ to infer the differentiation trajectory directionality and future cell state from ratio of un-spliced and spliced mRNAs within a single cell. Individual loom file was generated for each sample based on Cell Ranger output file using velocyto python package^115^. Then loom files were merged together for SLS samples and IS samples respectively. For visualization, we used Seurat generated single cell UMAP coordinates to project RNA velocity vectors onto the two-dimension embeddings.

#### Regulon and pathway analysis

Transcription factor enrichment and regulon activity were assessed using SCENIC package^45^ and human cisTarget databases: hg19-500bp-upstream-7species.mc9nr.feather and hg19-tss-centered-10kb-7species.mc9nr.feather. Seurat normalized expression matrix was used as input. Only the protein coding genes were analyzed for motif enrichment. We used clusterProfiler (v3.16.1)^116^ for pathway enrichment analysis with default parameters. The database used was Hallmark Gene Set from Molecular Signatures Database (MsigDB)^117^.

#### Visium spatial transcriptomics data processing

Raw FASTQ files were aligned to human GRCh38 reference genome using Space Ranger pipeline (10x Genomics). Raw data were processed using Seurat (v3.0.0) for normalization using SCTranform function. Custom scripts were used to map normalized spot-level data to histology images for visualization. Tumor enriched spots were identified as spots with averaged value of *CTSK* and *PMEL* higher than median of average value across all spots. We then calculated scores for SLS and IS cell states on tumor-enriched spots using the marker genes *MGP* (for SLS) and *ACTA2* (for IS) as these two marker genes were identified as most robust markers for these two distinct cell types in scRNA-Seq data. We used a stringent criterion to annotate SLS cell state: spots with value of *MGP* (SLS marker) higher than 75% across all spots and value of *ACTA2* (IS marker) lower than 75% across all spots. Vice versa, IS state was identified as spots with value of *ACTA2* (IS marker) higher than 75% across all spots and value of *MGP* (SLS marker) lower than 75% across all spots. SLS enriched island or IS enriched island were identified as island that only contain SLS state (blue) or IS state (green) tumor enriched spots. Since the expression levels of *TREM2* and *TYROBP* were within similar range, values of these two genes were simply averaged for each spot and plotted.

#### Spatial correlation analysis

For the major cell types identified in scRNA-seq analysis (Tumor cells, T cells, B cells, macrophages, lymphatic endothelial cells, blood endothelial cells, NK cells and dendritic cells), we re-defined cell type marker genes with the criteria that the relative average expression is 3 times higher than any other cell types and expressed in at least 50% of cells of the given cell type. The Nanostring spatial transcriptomics data were Q3 normalized and log2 transformed. Then, we calculated the relative frequency of each cell type in the Nanostring spatial transcriptomics datasets (12 bulk RNA-seq of ROIs of SLS tumor, and 12 bulk RNA-seq ROIs of IS tumor) by the average expression of scRNA-Seq re-defined cell type marker genes. To identify genes that may mediate cell-cell interactions, we performed Pearson correlation analysis of expression of genes that are primarily expressed in one cell type in the single cell data with the predicted frequency/activity of another cell type in the Nanostring spatial transcriptomics data of each ROI, followed by correlation test for significance assessment of the correlation coefficient. The assumption is that if a gene is highly expressed in one cell type and highly correlated with frequency/activity of another cell type, the given gene may mediate the interaction of these two cell types as previously described^90^.

#### Ligand-receptor interaction analysis

Ligand-receptor interaction analysis was performed to infer potential cell-cell interactions via direct ligand-receptor binding using algorithm described previously^86^. The set of ligand-receptor pairs were obtained from previous study^118^. We manually added more ligand-receptor pairs discovered more recently, including immune checkpoints and innate immune regulation. Briefly, interaction score of given ligand-receptor interaction between two cell types was calculated as the product of average ligand expression across all cells of one cell type and the average receptor expression across all cells of another cell types as previously described^86^. We calculated average expression of ligand and receptor in all cell types using normalized expression data of the aggregated scRNA-Seq dataset. The statistical significance of pairs of interaction was determined by one-sided Wilcoxon rank-sum test.

#### Kinetic modeling of purine metabolic pathway

As previously described^60^, we employed a kinetic model of purine metabolism^59^ that has the format of a Generalized Mass Action (GMA) system, where all processes are represented as products of power-law functions. The model contains 16 metabolites and 37 fluxes and a large number of regulatory signals^60^. The diagram of the model structure was drawn using custom scripts. We generated pseudo-bulk expression data from scRNA-Seq data by averaging expression of each gene across all non-zero cells in a given cell type. We used the differential expression of each gene in the tumor cells compared to matched normal mesenchymal cells as a corresponding change in enzyme amount. The enzyme activities were lumped into apparent rate constants in the original model formulation. Therefore, the differential expression of each gene was modeled as a corresponding change in its respective reaction rate constant parameter. All other parameters were retained the same as at the original steady state. The equations of the model were then integrated to get a new steady state where the variable concentrations and fluxes of the system were studied.

#### Reporting summary

Further information on research design is available in the Nature Research Reporting Summary linked to this article.

#### Data Availability

The scRNA-seq data of 5 LAM samples, 6 AML and 4 matched normal kidney samples that support the findings of this study are available in LAM Cell Atlas (https://research.cchmc.org/pbge/lunggens/LCA/LAM_projects.html) and GEO (data will be deposit before publication). All other data that support the findings of this study are provided in the supplementary files or available from the corresponding author upon request.

## SUPPLEMENTAL INFORMATION TITLES AND LEGENDS

**Supplementary Fig.1 Pathological images for AML and LAM specimen.**

**Supplementary Fig.2 Global single cell analysis of AML and LAM, related to** Fig. 1.

a. Expression of representative marker genes for cell types defined in AML. Red color shows the averaged normalized expression levels of marker genes.

b. Average expression of five tumor marker genes in AML (left) and LAM (right).

c. Graph-based clustering of mesenchymal cell population results in eight clusters.

d. Cell fraction for each AML patient across cell types.

e. Cell fraction for each LAM patient across cell types.

f. Expression of representative marker genes for macrophages and proliferating cells defined in LAM. Red color shows the averaged normalized expression levels of marker genes.

**Supplementary Fig.3 Marker gene expression in AML, related to** Fig. 1.

a. Identification of tumor cells (left) and tumor associated fibroblasts (TAF) (right) using marker genes shown. Cells expressed at least 2 marker genes above median value of corresponding genes across all cells were annotated as tumor cell or TAF, shown in red.

b. Upper panel: violin plots showing representative upregulated genes in AML cells. The y axis represents the normalized gene expression. Lower panel: feature plots of these genes.

c. Upper panel: violin plots showing expression of four long non-coding RNAs (lncRNAs). The y axis represents normalized gene expression. Lower panel: feature plots of these genes.

d. Representative regulons enriched in AML cells. First row: expression of transcription factors. Second row: regulon activities of these transcription factors.

e. Expression of *MDK* and *GPNMB* assessed by bulk RNA-Seq comparing AML tumors with normal kidneys.

f. Expression and regulon activity of *SP1* in AML cells.

g. Expression of *HIF1A* in AML cells.

**Supplementary Fig.4 Identification of two distinct cell states in AML/LAM cells, related to** Fig. 2.

(A) Feature plot showing averaged expression of *MGP*, *BGN*, *PRELP*, *LUM, MMP2, DCN* (left) and *TAGLN*, *CNN1*, *ACTA2*, *MYL9*, *MYH11*, *TPM2* (right).

(B) Feature plot showing expression of *TCF21*.

(C) Feature plot showing expression of *SOX4* and *TCF4*.

(D) Pathways enriched in SLS population, calculated using differentially expressed genes in cluster 1 (SLS) compared to cluster 2 (IS). Red: pathways enriched in SLS population; Blue: pathways enriched in IS population; x-axis: pathway activity score.

(E) Feature plots showing representative inflammatory genes upregulated in the IS population.

(F) Feature plots showing representative genes upregulated in the SLS population.

(G) Feature plots showing representative genes upregulated in the IS population.

(H) Expression of dormancy marker genes in TSC2-null 621-101 cells before and after estradiol treatment.

(I) RNAScope images of MDK expression in AML tumors.

**Supplementary Fig.5 Expression of marker genes in LAM, related to** Fig. 2.

(A) Feature plots showing expression of dormancy genes in LAM cells.

(B) ELISA assessment of serum MDK levels in healthy cohorts (n=19) and in LAM patients (n=20).

**Supplementary Fig.6 *In vitro* experiment showing SLS cells are rapamycin tolerant, related to** Fig. 3.

(A) Expression of genes identified as upregulated in AML cells in this study before and after rapamycin treatment in the primary culture.

(B) Expression of genes upregulated in SLS (refer to Supplementary Fig. 4f) in the control group.

(C) Expression of dormancy marker genes in the control group.

(D) Tumor volume relative to pre-treatment tumor volume. Tumor volume was measure immediately before each treatment. TTJ xenograft mice (n=6 per group) were treated 3 times/wk with DMSO, iMDK (9mg/kg), rapamycin (3mg/kg), or combined iMDK (9mg/kg) and rapamycin (3mg/kg).

(E) Expression of MDK across cancer types and matched normal tissues (data obtained from TCGA).

(F) Cell growth inhibited by combination treatment of rapamycin and iMDK on 3 bladder cancer cell lines. Left panel: microscopy pictures showing growth of 3 bladder cancer cell lines treated with 20nM rapamycin alone (first column) or combination treatment of 20nM rapamycin and 1µM iMKD (second column) for 6 days. Right panel: cell proliferation, assessed by crystal violet assay, of bladder cell line HT1376 on the treatment of DMSO (vehicle), 20nM rapamycin, 1µM iMDK or combination of 20nM rapamycin and 1µM iMDK for 14 days.

**Supplementary Fig.7 Remodeling of endothelial cells, related to** Fig. 4.

(A) Fraction of endothelial cells in IS and SLS tumors identified in IHC data.

(B) Expression of *VEGFA* in AML cells.

(C) Re-clustering of 20,772 endothelial cells (pooled from normal kidney and AML tumor) revealed 14 clusters.

(D) Expression of lymphatic endothelial marker genes (left) and blood endothelial marker genes (right) in endothelial cell clusters. One cluster of 646 cells (cluster 3; *PDPN*+, *PROX1*+ and *LYVE1+*) was composed of lymphatic endothelial cells (LECs) and was derived solely from tumor. Fourteen clusters of blood endothelial cells (*FLT1*+ and *PLVAP*+) were identified. Subtypes of endothelial cells were annotated based on published cell markers^119^.

(E) UMAP showing averaged expression of representative genes specifically expressed in endothelial cells (ECs) derived from AML tumors (left) or from matched normal tissues (right). Eleven of clusters were primarily tumor-derived (*PLVAP*+ and *APOLD1*+) and three clusters were primarily normal kidney-derived (*SOST*+ and *CRHBP*+).

(F) Feature plots showing expression of *CCL21* (C-C Motif Chemokine Ligand 21), *TBX1* and *NRP2*. These genes were observed specifically in tumor LECs. High expression of *CCL21* promotes immune cells migration and tumor metastasis^120^. High expression of *TBX1* is required for lymphatic vessel development^121^. NRP2 has been reported to be involved in sprouting lymphangiogenesis^122^.

(G) Hallmark pathways enriched in tumor-associated endothelial cells. Analysis of hallmark pathway gene signatures comparing all tumor-derived endothelial cells versus normal kidney-derived endothelial cells revealed stronger fatty acid metabolism in tumor-associated endothelial cells. Fatty acid metabolism plays a critical role in lymphatic differentiation and development by providing energy resources for cell proliferation and epigenetic regulation^123^.

(H) Expression and regulon activity of *NR2F1* and *NR2F2*. Left two panels: expression of *NR2F1* and *NR2F2;* right two panels: regulon activity of these transcription factors. Regulon analysis identified candidates that may underlie these gene expression differences in LECs, including transcription factors such as *NR2F1* and *NR2F2* that are highly expressed in tumor-derived LECs specifically. The genetic networks regulated by these transcription factors also showed high activity in LECs. NR2F2 is of particular interest in LAM because of its identification as a GWAS susceptibility locus^124^. NR2F2 has been shown to physically and functionally interact with PROX1 to regulate LEC-fate specification^125, 126^ and promote the formation of lymphatic vasculature^122^. NR2F2 controls sprouting lymphangiogenesis by transcriptional up-regulation of NRP2, a co-receptor for VEGF-C^127^. NR2F1 has recently been shown to be required for vascular development^128^.

**Supplementary Fig.8 Characterization of T cells related to** Fig. 5.

(A) Pathways enriched in T cells from tumor samples compared to those from matched normal tissues. Red bars represent pathways enriched in T cells from tumor samples; blue bars represent pathways enriched in T cells from matched normal tissues.

(B) UMAP plot of cell origin.

(C) Averaged expression of *MKI67*, *MCM7*, and *PCNA* in T cells.

(D) CD3 IHC on a representative AML tumor and matched normal kidney.

(E) Feature plots showing expression of checkpoint markers in T cells.

(F) Bar plot showing fraction of T cells in SLS versus IS dominant tumors.

(G) Expression profiles of immune checkpoint ligands: TIGIT ligands (*PVR*, *NECTIN2*), BTLA ligand (*TNFRSF14*), LAG3 ligand (*HLA-DRA*, *FGL1*), *KLRG1* ligand (*CDH1*, *CDH2*), and PD-1 ligands (*CD274*, *PDCD1LG2*). The y axis represents the normalized gene expression value.

**Supplementary Fig.9 Suppressive immune microenvironment in AML, related to** Fig. 6.

(A) Violin plot of expression of immune checkpoint genes in macrophages obtained from tumors or from matched normal kidneys.

(B) Images of Nanostring ROIs.

(C) Inferred interactions between tumor cells and macrophages calculated by integrative analysis of spatial transcriptomics of the representative IS-dominant tumor (12 ROIs) and scRNA-Seq. x-axis displays relative expression of genes in single cell data. Only genes that are expressed in both single cell data and spatial transcriptomics data are shown. Left side are genes relatively highly expressed in tumor cells; right side are genes relatively highly expressed in macrophages. Y-axis displays Pearson Correlation Coefficient (PCC) of gene expression with macrophage frequency in spatial transcriptomics data. Genes with log-ratio less than −1.5 and correlation coefficient higher than 0.4 are colored. *APOE*: PCC=0.49, p=0.1 (correlation test).

(D) Similar analysis as in D calculated by integrative analysis of bulk RNA-seq data from a published dataset^8^ and scRNA-seq. *APOE*: PCC=0.55, p=0.1 (correlation test).

(E) Feature plots showing expression of *IL7R*, *GZMK*, *GZMH*, *IFITM1* in macrophages.

(F) Feature plots showing expression of marker genes of tumor-associated macrophages, including *CD163*, *MRC1*, *TREM2*, *VEGFA*.

**Supplementary Fig.10 Molecular interactions between tumor and tumor microenvironment inferred by ligand-receptor co-expression, related to** Fig. 7.

(A) Pathways enriched in plasma cells and follicular B cells obtained from tumors, and enriched in B cells obtained from matched normal tissues.

(B) Feature plots showing expression of *CLEC9A* and *XCR1* in dendritic cells.

(C) Pathways enriched in each cluster of dendritic cells colored by column bar.

(D) Interactions between B cells and other cell types in SLS-dominant tumors, calculated as the product of the average ligand expression and average receptor expression. Only interactions with a score greater than 1 across any cell type pair are displayed. Each column shows a pair of cell types, and each row shows the ligand-receptor pair. The color indicates interaction score. Column label: cell type expressing the ligand and cell type expressing the receptor are separated by “_”. Row label: ligand and receptor are separated by “_”.

(E) Interactions between T cells and other cell types in SLS-dominant tumors, calculated as the product of the average ligand expression and average receptor expression. Only interactions with a score greater than 1 across any cell type pair are displayed.

(F) Interactions between NK cells and other cell types in SLS-dominant tumors. Only interactions with a score greater than 1 across any cell type pair are displayed.

(G) Interactions between tumor cells and other cell types in SLS-dominant tumors. Only interactions with a score greater than 1 across any cell type pair are displayed.

**Supplementary data 1. Patient information.**

**Supplementary data 2. Differentially expressed genes in Tumor, TAF and normal mesenchymal cells, related to** Fig. 1g**. Each of cell type was compared to the other two cell types separately.**

**Supplementary data 3. Enriched regulons in SLS tumor cells compared to matched normal mesenchymal cells, related to** Fig. 1l.

**Supplementary data 4. Enriched regulons in IS tumor cells compared to matched normal mesenchymal cells, related to** Fig. 1l.

**Supplementary data 5. Marker genes of each cluster of primary culture cells.**

**Supplementary data 6. Re-defined gene signatures for each major cell type based on scRNA-Seq data.**

**Supplementary data 7. Identified genes mediating cell-cell interactions, related to** Fig. 6d.

**Supplementary data 8. Molecular interactions identified between TAF and SLS or IS tumor cells.**

